# Global Motion Detection and Censoring in High-Density Diffuse Optical Tomography

**DOI:** 10.1101/2020.02.22.961219

**Authors:** Arefeh Sherafati, Abraham Z. Snyder, Adam T. Eggebrecht, Karla M. Bergonzi, Tracy M. Burns-Yocum, Heather M. Lugar, Silvina L. Ferradal, Amy Robichaux-Viehoever, Christopher D. Smyser, Ben J. Palanca, Tamara Hershey, Joseph P. Culver

## Abstract

Motion-induced artifacts can significantly corrupt optical neuroimaging, as in most neuroimaging modalities. For high-density diffuse optical tomography (HD-DOT) with hundreds to thousands of source-detector pair measurements, motion detection methods are underdeveloped relative to both functional magnetic resonance imaging (fMRI) and standard functional near-infrared spectroscopy (fNIRS). This limitation restricts the application of HD-DOT in many challenging situations and subject populations (e.g., bedside monitoring and children). Here, we evaluate a new motion detection method for multichannel optical imaging systems that leverages spatial patterns across channels. Specifically, we introduce a global variance of temporal derivatives (GVTD) metric as a motion detection index. We show that GVTD strongly correlates with external measures of motion and has high sensitivity and specificity to instructed motion - with area under the receiver operator characteristic curve of 0.88, calculated based on five different types of instructed motion. Additionally, we show that applying GVTD-based motion censoring on both task and resting state HD-DOT data with natural head motion results in an improved spatial similarity to fMRI mapping for the same respective protocols (task or rest). We then compare the GVTD similarity scores with several commonly used motion correction methods described in the fNIRS literature, including correlation-based signal improvement (CBSI), temporal derivative distribution repair (TDDR), wavelet filtering, and targeted principal component analysis (tPCA). We find that GVTD motion censoring outperforms other methods and results in spatial maps more similar to matched fMRI data.

## 1 Introduction

High-density diffuse optical tomography (HD-DOT) has tremendous potential to be a surrogate for functional magnetic resonance imaging (fMRI) [1-6]. However, methods for dealing with detection and suppression of motion artifacts in HD-DOT data are relatively underdeveloped, which limits its application to many important clinical populations. While fMRI has become a gold standard for cognitive neuroimaging, it is contraindicated in subjects with metal implants and cannot be used in many clinical settings, and studies seeking more naturalistic imaging environments. In contrast, fNIRS-based methods are portable, suitable for naturalistic imaging, and not contraindicated in subjects with electronic or metal implants [7-17]. Sparse fNIRS imaging arrays yield poor resolution and low image quality. HD-DOT provides improved image resolution and depth profiling, particularly when used with anatomical head models [18-20]. However, as in both fMRI and fNIRS, detection, classification, and removal of motion-induced artifacts remains a challenge for HD-DOT.

Multiple fMRI studies have documented the spurious effects of motion artifacts in blood oxygen level-dependent (BOLD) fMRI despite the use of common motion suppression methods [21-26]. Motion-induced changes in T2*-weighed fMRI signals are shared across brain voxels, hence generate spatially structured artifacts. Such artifacts alter functional connectivity by decreasing long-distance correlations and increasing short-distance correlations [21, 24-26]. However, two simple data quality indices, frame-wise displacement (FD) and root mean squared (RMS) signal change over sequential frames (DVARS), are commonly used in fMRI data processing pipelines to identify and exclude data segments (motion censoring or scrubbing) from behaviorally relevant fMRI measures [21, 27, 28].

In HD-DOT, similar to fMRI, the effects of head motion are global across the field of view (FOV) and impact a majority of measurements or voxels. In fMRI, head movements shift the position of the brain in space and modulate the BOLD signal [29, 30], in HD-DOT, head motion induces a torque on the fibers in the optical imaging array that, in turn, modulates the location (Fig. 1B center), angle, or both location and angle of optode-scalp coupling (Fig. 1B right). Thus, motion induces artifacts in the optical signals that can appear as brief transient spikes or baseline shifts. These artifacts propagate from measurement space to voxel space in the image reconstruction process and corrupt the neuroimaging results.

**Figure 1.**
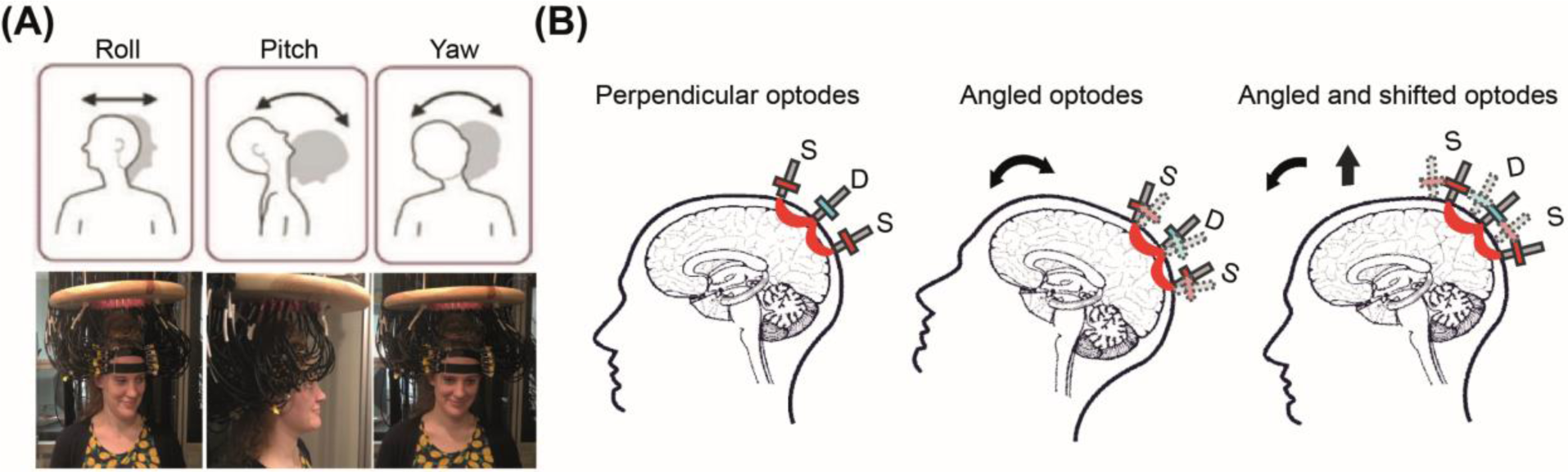
Effects of head motion on HD-DOT optode coupling. (A) Research participant wearing an HD-DOT imaging cap. Head rotation may occur about three axes (roll, pitch, and yaw). (B) Schematic illustration of how head motion can affect optode couplings. The far-left figure shows the ideal perpendicular angle between the SD optodes and the head. The middle figure shows the angled optodes as a result of nodding up and back to center. The far-right figure shows the angled and shifted optodes as a result of nodding up and body movement.

Numerous strategies for managing motion-induced artifact have been described in the fNIRS literature. However, a consensus on how best to correct for motion artifacts has not emerged [31-33]. Extant motion correction methods in fNIRS largely involve two steps: first, motion detection, and second, signal correction [34-39]. The fNIRS literature has largely focused on correcting motion artifacts on individual source-detector pair measurements and much less attention has been placed on multichannel or full arrays assessments. Moreover, most fNIRS studies have not assessed the efficacy of the denoising methods through comparison against fMRI.

We address these limitations by conducting a comprehensive evaluation of motion artifact removal methods for HD-DOT data by including independent measures of motion (accelerometry) and comparisons against gold-standard matched fMRI datasets. We introduce a novel index of motion, the global variance of the temporal derivatives (GVTD), and show that it strongly correlates with directly transduced measures of motion and outperforms two commonly used temporal motion detection indices in fNIRS based on the single-channel changes in the signal amplitude. We then optimize the use of GVTD-based motion detection in the HD-DOT processing pipeline by measuring the artifact-to-background ratio in in vivo resting state datasets collected with different HD-DOT devices in adults and infants. Using GVTD as a quantitative index, we show that GVTD predicts the quality of the task-based brain response maps (where the quality is defined based on the voxel-wise similarity between the HD-DOT images and matched fMRI data). Finally, we investigate the efficiency of the GVTD-based motion detection and censoring on the HD-DOT task and resting state datasets from exemplar HD-DOT datasets. These analyses demonstrate that GVTD censoring outperforms current fNIRS motion correction methods.

## 2 Methods

### 2.1 The global variance of the temporal derivatives (GVTD)

GVTD indexes global instantaneous change in the optical time-courses. For each time point, GVTD is computed as the RMS of the temporal derivatives across a set of measurements (Eq. 1). In this paper, the first nearest neighbor measurements (nn1) with a source-detector (SD) distance of 13 mm (10 mm for infants) were chosen, as they are more sensitive to changes in the fiber-scalp coupling and relatively insensitive to brain dynamics in comparison to longer distance measures [40].

The simple analytic formula for GVTD is

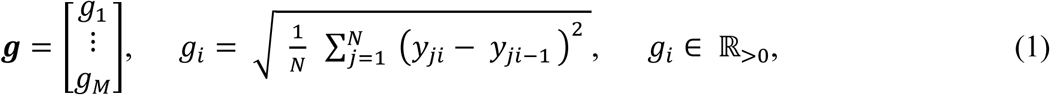

where ***g*** is the GVTD vector, *y*_*ji*_ *ϵ* ℝ is the optical density change or molar HbO_2_ at spatial coordinate *j. i* indexes the time points, *N* is the number of coordinates, and *M* is the number of time-points.

### 2.2 Motion censoring in HD-DOT data

Motion censoring (scrubbing) excludes the time-points (blocks) exceeding the GVTD noise threshold from further analysis of resting state and task data [41, 42]. Details concerning the noise threshold criterion are explained in §3.3. This proposed HD-DOT censoring strategy follows a similar practice that resulted in statistical improvements in the resting state as well as task fMRI data [21, 43-45].

### 2.3 Other comparison motion correction methods

#### 2.3.1 Correlation-based signal improvement (CBSI)

CBSI motion correction is based on the assumption that oxygenated and deoxygenated hemoglobin signals are negatively correlated under all circumstances. In the presence of motion artifacts, the correlation between these two signals becomes more positive. CBSI corrects the oxyhemoglobin concentrations by subtracting the scaled deoxy-hemoglobin to match the variance of the oxygenated signal. This process removes the positive correlation content between the two signals, taking into account their different amplitudes. Then, the corrected deoxy-hemoglobin is calculated by multiplying the corrected oxy-hemoglobin by the inverse of the same scaling factor between the original signals [37]. In this paper, we performed this motion correction method after spectroscopy on the down-sampled 1 Hz (Fig. S1).

#### 2.3.2 Targeted principal component analysis (tPCA)

Principal component analysis (PCA) projects an arbitrary set of signals onto orthogonal principal components. Then, the principal components with the least variance are excluded, and the signal is reconstructed from the remaining components. Targeted PCA (tPCA) applies PCA to temporal epochs of the data that is identified to contain motion artifacts. tPCA reduces the risk of eliminating the physiological content in the motion-free epochs of the signal [46]. Hence, this method is followed by a prior step of motion detection in the temporal domain. Conventionally, this motion detection is performed by setting a threshold on signal amplitudes or the windowed signal amplitude changes. In this paper, we used the Homer function “hmrMotionArtifactByChannel” to detect noisy timepoints and “hmrMotionCorrectPCA” to perform PCA, and set the parameters of this algorithm in the similar range as in the original study [46]; tMotion = 0.5, tMask = 2, STDEVthresh = 20, AMPthresh = 0.5, nSV = 0.97 (Tables S1 and S2, Fig. S1).

#### 2.3.3 Wavelet filtering

Wavelet-based motion correction is based on discrete wavelet transformation of single-channel measurements. This method assumes that the distribution of the wavelet coefficients of a motion-free signal should follow a Gaussian distribution. Therefore, motion artifacts are detected based on the deviations from the Gaussian distribution. By setting an outlier detection threshold, the coefficients associated with motion artifacts are excluded, and the clean signal is reconstructed based on the remaining wavelet coefficients [38]. We used the “hmrMotionCorrectWavelet” function, setting the interquartile parameter as 1.5, as suggested in the original paper [38] (Tables S1 and 2S, Fig. S1).

#### 2.3.4 Kurtosis-based wavelet filtering (kbWF)

The kurtosis-based wavelet filtering (kbWF) method optimizes the use of the wavelet filtering motion correction by setting the threshold based on the kurtosis of the coefficient distributions [39]. The “*hmrMotionCorrectKurtosisWavelet”* function was used with the kurtosis threshold parameter set to 3.3, as recommended in the original paper [39] (Tables S1 and 2S, Fig. S1).

#### 2.3.5 Hybrid (Spline + Savitzky Golay)

The spline and Savitzky-Golay hybrid method is a three-step algorithm that aims to identify and correct different types of motion artifact [36]. First, single-channel measurements are passed through a Sobel filter to identify time-points exceeding a threshold of 1.5 times the interquartile interval of the signal gradient. Second, this method performs a spline interpolation on those epochs containing motion to remove the baseline shifts and slow spikes. Steps 1 and 2 were introduced in a previous fNIRS motion removal method, commonly known as the motion artifact removal algorithm (MARA) [34]. After this step, the hybrid method then applies a Savitsky-Golay smoothing filter to remove the remaining fast spikes. We used the “*hmrMotionCorrectSplineSG”* function defined in the original paper with its default parameters and setting p = .99 and FrameSize_sec = 1.5 [36] (Tables 1S and 2S, Fig. S1).

#### 2.3.6 Temporal derivative distribution repair (TDDR)

Temporal Derivative Distribution Repair (TDDR) also is a three-step algorithm that aims to automatically identify and correct motion artifacts at the single-channel level. First, by computing the temporal derivative of the signal, TDDR initializes the vector of observation weights. Second, it iteratively estimates the robust observation weights by applying the resulting robust weights to the centered temporal derivative to produce the corrected derivative. Finally, it integrates the corrected temporal derivative to yield the corrected signal [35].

### 2.4 Independent measurement of head motion

A motion sensor (3-space™ USB/RS232; Yost Labs, Portsmouth, Ohio) was attached to the top strap of the HD-DOT cap in a subset of the data acquired with instructed motion (more details in 2.6.2). This sensor includes a triaxial inertial measurement unit (IMU), which uses a gyroscope, an accelerometer, and a compass sensor (Fig. S2). Onboard electronics compute and report in real-time, the quaternion-based orientation relative to an absolute reference. We synchronized the outputs of the motion sensor with our HD-DOT data acquisition system using audio pulses at the start and end of data streams. The motion sensor data were down-sampled from 200 Hz to 1 Hz to match the final sampling rate of the HD-DOT data. Then, the motion sensor and HD-DOT signals were aligned by delaying the earlier signal based on the cross-correlation delay time with maximum correlation value.

### 2.5 Angular rotation

The angular rotation (**Φ**) time-course was defined as the norm of the temporal derivatives of the head orientation in terms of Euler angles (*α* roll, *β* pitch, and *γ* yaw), measured by the motion sensor. This index was defined in a manner similar to that of GVTD to facilitate comparisons between GVTD and motion sensor outputs (Eq. 2).

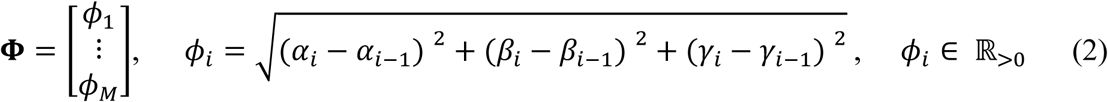

In this notation, *i* indexes the time points, and *M* is the number of time-points.

### 2.6 Artifact-to-background ratio (ABR)

To quantify the magnitude of the motion artifacts, we defined the artifact-to-background ratio (ABR; *ρ*), where ABR is the mean GVTD of all time-points above the noise threshold (defined in §3.3), divided by the mean GVTD of all the time-points below the noise threshold (Eq. 3).

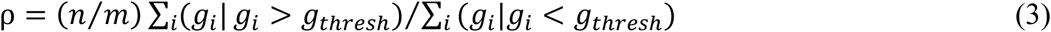

In this formula, *g*_*i*_ is the GVTD value at time index *i, g*_*thresh*_ is the threshold value, *n* is the number of time-points below the threshold, and *m* is the number of time-points above the threshold.

### 2.7 Datasets and general data processing

#### 2.7.1 Datasets and their objective

Dataset 1: For validation, we collected an fMRI dataset in which adult subjects (n = 8) were scanned in both the resting state and during a hearing words (HW) task. This dataset served as ground truth. Dataset 2: As a positive control, in this HD-DOT dataset, healthy adults (n = 12) performed instructed motion while performing the same HW task performed during fMRI. Dataset 3: In this HD-DOT dataset, adult subjects (n = 13) performed the same HW task without instructed motion. Dataset 4: In this HD-DOT dataset, healthy adults (n = 8) were scanned while awake in a task-free (resting) state. Dataset 5: In this HD-DOT dataset, healthy term infants (n = 11) were imaged in the resting state (awake or asleep). This is a previously published dataset [3]. Demographic information and the objective of using each dataset are reported in Table 1.

**Table 1:**
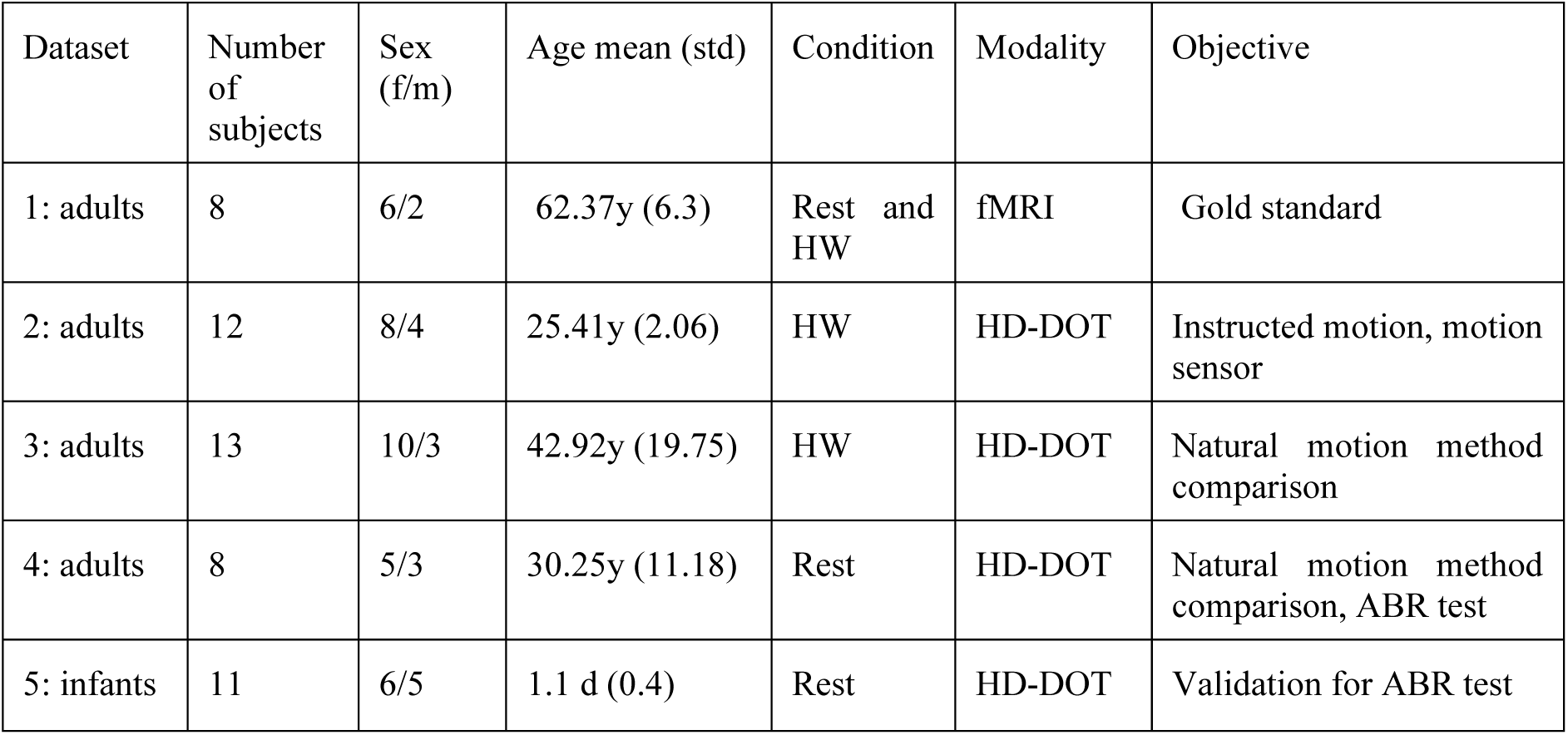
Demographic information. HW: hearing words; y: year; d: day; ABR: artifact-to-background ratio.

| Dataset | Number of subjects | Sex (f/m) | Age mean (std) | Condition | Modality | Objective |
| --- | --- | --- | --- | --- | --- | --- |
| 1: adults | 8 | 6/2 | 62.37y (6.3) | Rest and HW | fMRI | Gold standard |
| 2: adults | 12 | 8/4 | 25.41y (2.06) | HW | HD-DOT | Instructed motion, motion sensor |
| 3: adults | 13 | 10/3 | 42.92y (19.75) | HW | HD-DOT | Natural motion method comparison |
| 4: adults | 8 | 5/3 | 30.25y (11.18) | Rest | HD-DOT | Natural motion method comparison, ABR test |
| 5: infants | 11 | 6/5 | 1.1 d (0.4) | Rest | HD-DOT | Validation for ABR test |

All aspects of these studies were approved by the Human Research Protection Office of the Washington University School of Medicine. All adult participants in the previous and new datasets were right-handed, native English speakers, and reported no history of neurological or psychiatric disorders. Adults were recruited from the Washington University campus and the surrounding community (IRB 201101896, IRB 201609028). All full-term infants were recruited from the Newborn Nursery at Barnes-Jewish Hospital in St Louis, Missouri, within the first 48 hours of life (IRB 201101813). All subjects (or their guardians) gave informed consent and were compensated for their participation in accordance with institutional and national guidelines.

#### 2.7.2 HD-DOT systems, image reconstruction, and spectroscopy

All adult HD-DOT datasets (datasets 2, 3, and 4) were collected using a previously described continuous-wave HD-DOT system comprising 96 sources (LEDs, at both 750 and 850 nm) and 92 detectors (coupled to avalanche photodiodes, APDs) [1]. Acquisition in infants was performed at the bedside using a previously reported portable continuous-wave HD-DOT system with an optode array consisting of 32 sources (LEDs, at both 750 and 850 nm) and 34 detectors [3]. More detailed descriptions of the imaging systems are given in the corresponding references. Light modeling was computed using the standard MNI atlas-based absorption model; details can be found in [18]. Volumetric movies of relative changes in absorption at 750 nm and 850 nm were reconstructed after inverting the sensitivity matrix using Tikhonov regularization and spatially variant regularization [1]. Relative changes in hemoglobin concentration were obtained via a spectral decomposition of the absorption data, as previously described [1, 3].

#### 2.7.3 Functional MRI (fMRI) system and imaging

All fMRI data were collected on a research-dedicated Siemens 3.0T Magnetom Prisma system (Siemens Medical Solutions, Erlangen, Germany), with an iPAT compatible 20-channel head coil. Blood Oxygenation Level Dependent (BOLD) sensitized fMRI data with TR = 1230 ms, TE = 33 ms, voxel resolution = 2.4 mm^3^, FA = 63 degrees, with a multi-band factor of 4 for both resting state functional connectivity MRI (3 runs each 10 min) and HW task BOLD (1 run, 3.5 min) were acquired for all subjects in dataset 1.

### 2.8 Paradigms

Hearing words: Subjects were seated for HD-DOT or supine for fMRI and instructed to fixate on a white crosshair against a gray background while listening to words. The HW task was administered as a block design. Each trial consisted of 15 seconds of hearing words followed by 15 seconds of silence. Each run included multiple trials, n = 10 for dataset 2, and n = 6 for datasets 1 and 3. The total number of acquired runs per session was 7 (dataset 2) or 1 (datasets 1 and 3).

Instructed motion: The instructed motion was performed by subjects during the HW task (dataset 2), with 15% of the trials including instructed motion. Participants viewed a screen with a crosshair and were instructed to perform a specific motion type when the crosshair color changed. Movements were performed for about 2 seconds every 3-5 seconds over a 15-second word presentation section. Subjects were monitored in real-time using a digital camera to ensure that they were engaged in the assigned tasks. Specific motions included (i) head turn to the left and back to center (roll, Fig. 1A left), (ii) head nod up and back to center (pitch, Fig. 1A center), (iii) shifting body position, (iv) taking deep breaths, and (v) raising eyebrows. Head twist (yaw, Fig. 1A right) motion was avoided to prevent cap displacement.

Resting state: Resting state data in adults (datasets 1 and 4) were collected over 10 min runs while subjects were seated for HD-DOT or supine for fMRI and visually fixating on a white crosshair against a gray background. Subjects were asked to stay awake and still during data acquisition. The number of runs per session was 3 (dataset 1) or 1 (datasets 4). Resting state HD-DOT in infants was acquired at the bedside (dataset 5) within the first 24-48 hours of life during natural (un-medicated) sleep or quiet rest [3].

### 2.9 Data processing

#### 2.9.1 HD-DOT pre-processing

All HD-DOT data were processed using the NeuroDOT toolbox following the flowchart in Fig. S1 [1, 47, 48]. HD-DOT light measurement data were converted to log-ratio (using the temporal mean of a given SD-pair measurement as the relative baseline for that measurement). Noisy measurements were empirically defined as those with greater than 7.5% temporal standard deviation in the least noisy (lowest mean GVTD) 60 seconds of each run [19], and were excluded from further processing. Then the data were high-pass filtered (0.02 Hz cut-off for task-based datasets, 0.009 Hz for resting state datasets) to remove low-frequency drift. To serve as an estimate of the global superficial signal, we computed the average of all remaining first nearest neighbor measurements (13 mm SD-pair separation in the adult system and 10 mm SD-pair separation in the infant system). This global signal estimate was regressed from all measurements [40]. After low-pass filtering (0.5 Hz cut-off for task-based data sets, 0.08 Hz for resting-state data sets), the time-courses were down-sampled from 10 Hz to 1 Hz and then used for image reconstruction. The efficacy of GVTD was evaluated at four stages of the HD-DOT processing pipeline, as indicated in Fig. S1 (green boxes) on 10 Hz sampled data. All other motion correction methods except CBSI were also performed on the 10 Hz sampled optical density signals (immediately after the log-ratio step) (Fig. S1).

#### 2.9.2 fMRI pre-processing

fMRI pre-processing was performed using in-house 4dfp tools [49]: 1. correction for systematic slice-dependent time shifts; 2. elimination of odd-even slice intensity differences due to interleaved acquisition; 3. rigid-body realignment for head motion within and across runs; 4. normalization of signal intensity to a mode value of 1000. Signal intensity normalization enables identification of artifact by evaluation of the signal temporal derivative. Atlas transformation was computed by composition of affine transforms derived by a sequence of coregistration of the fMRI volumes via the T2-weighted and MP-RAGE structural scans. Head motion correction and atlas transformation was applied in a single resampling step that generated volumetric time series in (3mm)^3^ atlas space. Data underwent spatial smoothing (6 mm full width at half maximum in each cardinal direction) and temporal band-pass filtering (0.02-0.5 Hz for the HW task and 0.009-0.08 for resting state). Nuisance regressors included six rigid body values derived from head motion correction, white matter, and CSF signals, and the mean whole-brain signal. Motion artifacts were reduced in resting state data through DVARS-based motion scrubbing using session-specific thresholding expressible as 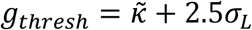 (see Eq. 5 below) [50]. The fraction of censored frames was 21% ± 12%.

#### 2.9.3 HW task response mapping in datasets 1, 2, and 3

Another objective of acquiring HW task data was to evaluate GVTD as an index of HD-DOT data quality (dataset 2). To this end, 70 trials of HW (15 sec of HW (On), 15 sec of silence (Off)) were acquired in each session; 10 trials included instructed motion; the remaining 60 trials (ordinary trials) did not. The reconstructed voxel-wise data represent the changes in the hemoglobin concentration (*Δ*[*HbO*_2_]) in units of *μ*mol/L [51]. The quantitative response magnitude was then calculated with a standard general linear model (GLM). The design matrix was constructed by convolving the experimental design with a canonical HRF using a two-gamma function fitted to the in-vivo HD-DOT data, as described in [52]. Extracted hemodynamic response estimations for each subject were then combined in a simple group-level fixed effects analysis [53]. Fixed effect analysis was adopted as we expect the variance in our dataset to be most strongly driven by scan-to-scan variability rather than from subject-to-subject differences.

#### 2.9.4 Seed-based correlation analysis of functional connectivity in datasets 1 and 4

Seed regions were 5 mm radius spheres centered on coordinates used in our previous study [1]. Five seeds representing the auditory (AUD), visual (VIS), somatomotor (MOT), dorsal attention network (DAN), and frontoparietal network (FPN) networks were selected within the HD-DOT FOV. Correlation maps were generated by calculating the Pearson correlation between the time-series of each seed region with all other voxels in the FOV. Correlation maps in individuals were Fisher’s z-transformed and averaged across subjects.

## 3 Results

### 3.1 Effect of motion artifacts on HD-DOT data

We investigated the effects of various types of movements on HD-DOT data using instructed motion. During the HW task, subjects performed five different types of instructed motion including large movements (head rotation) and small movements (raising eyebrows) (§2.8). One way to track the effect of motion is to spatially display the measurement pair channels (Fig. 2B). For example, for all the second nearest neighbor (nn2) pairs, we can mark sources and detectors with very high standard deviations over time during instances of instructed roll rotation (pink circles) and eyebrow motion (blue circles) (Fig. 2B). Alternatively, one can analyze an SD-pair measurement (pair highlighted by large circles in Fig. 2B) by comparing its time-course during runs without instructed motion (“ordinary”), or with different levels of instructed motion, i.e., low eyebrow motion or gross roll rotation (Fig. 2C). The difference in signal quality between the clean and corrupted responses are evident after block averaging (Fig. 2D).

**Figure 2.**
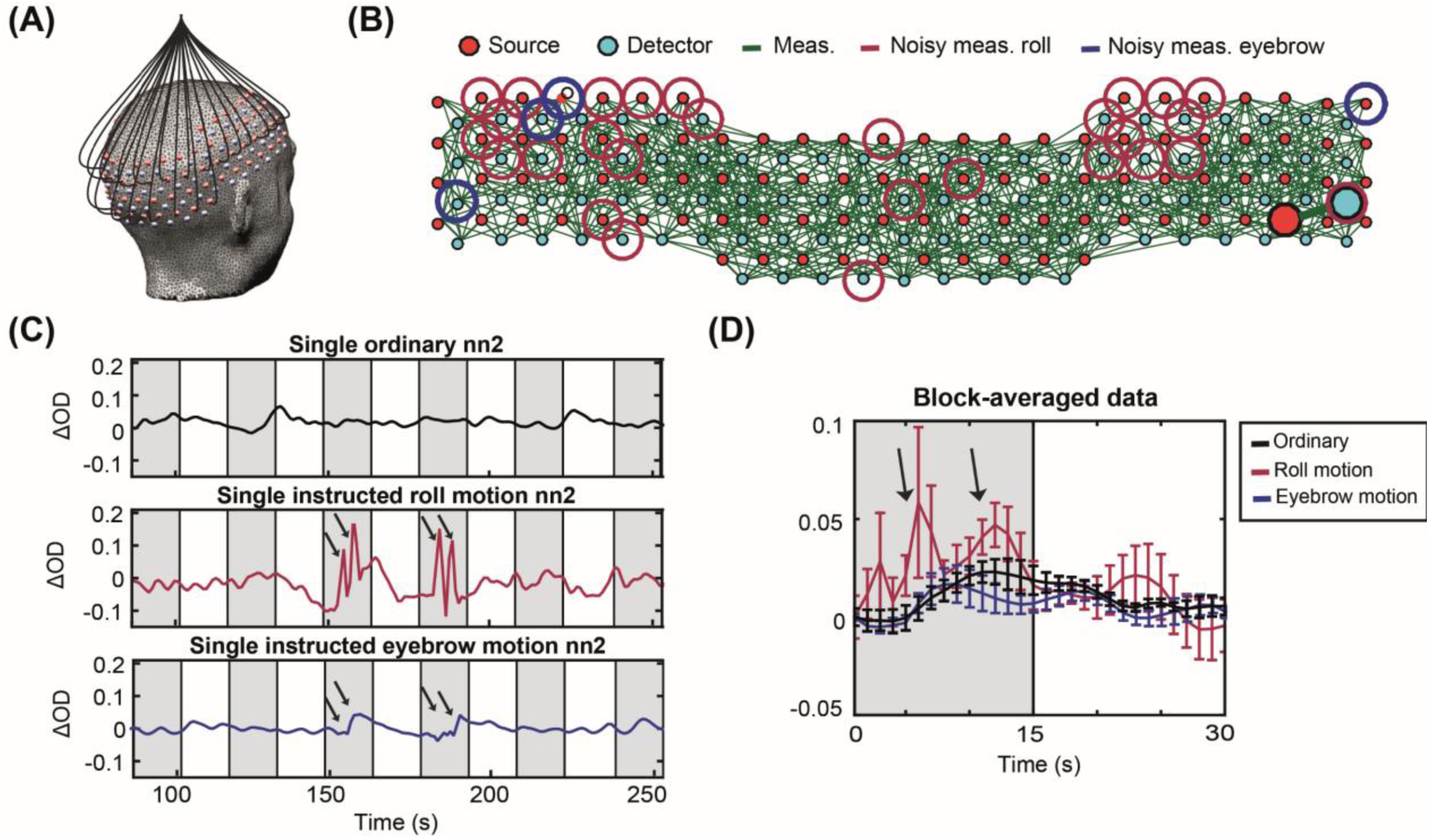
(A) Adult HD-DOT cap structure illustrating a subset of optical fibers. (B) Green lines indicate source-detector (SD) pairs that have a standard deviation less than 7.5%. Source or detector locations identified as noisy for roll (large red circles) and eyebrow (large blue circles) motion, respectively. (C) Changes in the light levels of a representative SD-pair during HW runs that were ordinary (black), instructed roll motion (pink), and instructed eyebrow motion (blue). Arrows indicate motion. Gray shading indicates auditory stimulus presentation. (D) Block averages of ordinary (black), instructed roll motion (pink) or instructed eyebrow motion run (blue). Error bars represent the standard error of the mean across trials.

We assessed the effects of different motion artifacts on the measurements by calculating the number of measurements with excessive noise for each type of motion artifact across all subjects. The HD-DOT array contains n = 1500 total measurements per wavelength within nn1 ∼ 13 mm, nn2 ∼ 30 mm, nn3 ∼ 39 mm, and nn4 ∼ 47 mm separations, respectively. All five motion types affected multiple SD channels distributed across the FOV; specifically, 51 ± 8% of the channels for gross body movement and 39 ± 4 % for small eyebrow movement. Based on these observations, we concluded that each type of motion generates global effects. Therefore, we adopted the GVTD as a global index of motion, taking into account optical signals over the full FOV.

### 3.2 GVTD and its correlation with the head angular rotation

The global effect of motion artifacts in HD-DOT can be visualized as a matrix where each row is a measurement signal and the columns index time (Fig. 3A). This type of visualization is similar to fMRI “gray plots” [21, 54, 55]. Inspection of Fig. 3A reinforces the notion that the effects of head motion in HD-DOT are global. GVTD time-course is computed in four steps. First, starting from the matrix of 850 nm nn1 optical density changes (Fig. 3A), the matrix of the backward differentiation of the selected time-courses is calculated (Fig. 3B). Then, from the matrix of the squares of backward differences (Fig. 3C), GVTD is defined as the square root of the mean across the selected measurement array (Fig. 3D). This sequence of steps progressively increases the sensitivity and specificity of the measure to motion (Fig. 3A-D).

**Figure 3.**
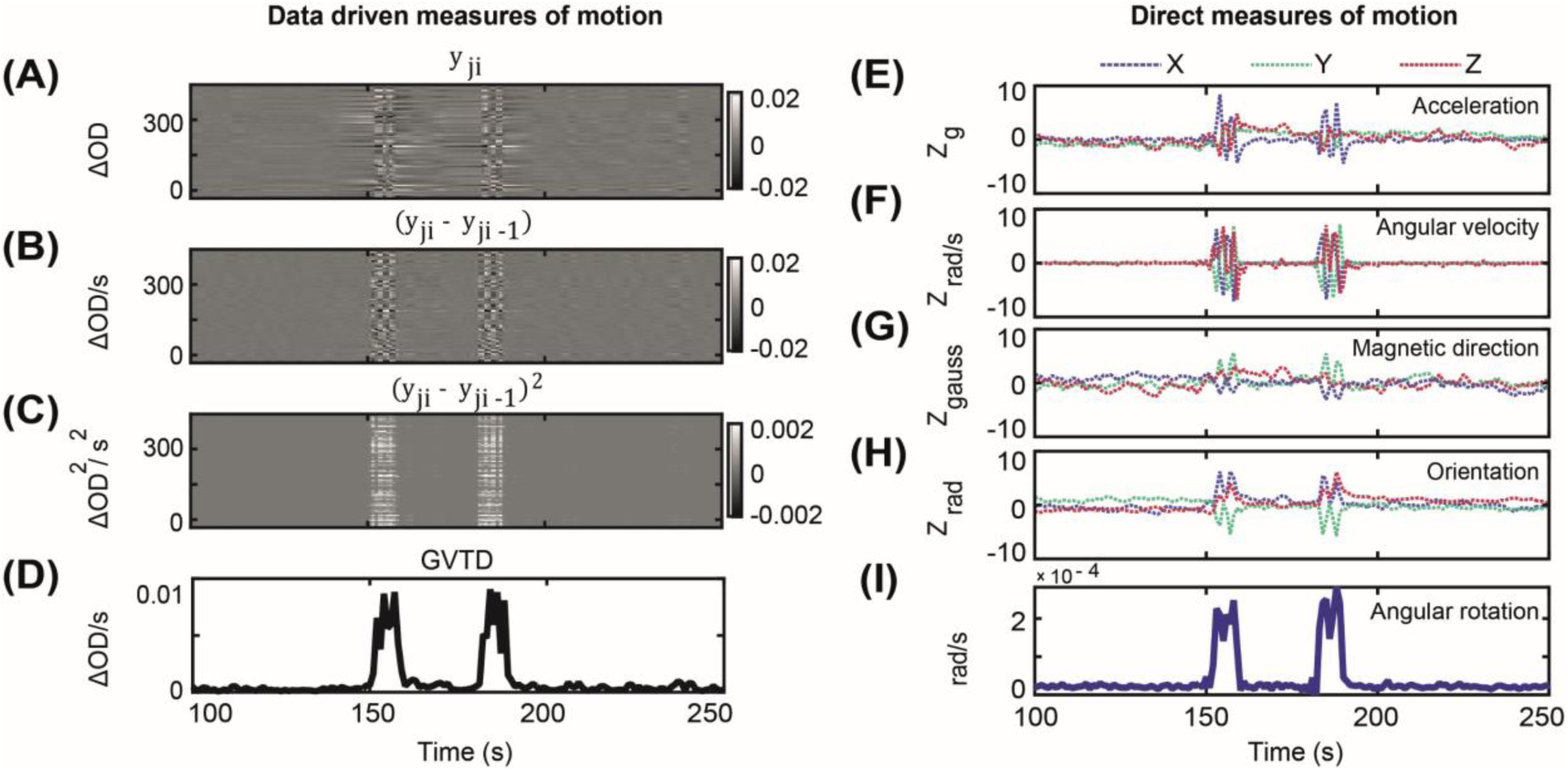
(A) All 850 nm nn1 measurements (n = 322) for a run containing instructed roll motion, represented as a matrix of measurements by time. (B) Temporal derivative of the data shown in (A); note intensified contrast between instructed motion vs. neighboring time-points. (C) Squared values (by element) of the matrix shown in (B). (D) GVTD time-course calculated as the RMS of the square values shown in (C). (E-H) Standardized (Z-scored) time-courses captured during instructed head motion in one subject. Colored traces correspond to x-, y-, and z-axes of the (E) accelerometer, (F) gyroscope, (G) compass, and (H) head orientation. (I) Angular rotation is calculated as the norm of the temporally differentiated x, y, and z time-courses shown in (H).

To evaluate the sensitivity of GVTD to motion, we concurrently recorded accelerometry as an independent measure in a subset of our instructed motion dataset (Fig. 3E-H). The graded quantitative motion capture of the accelerometer provided insight into the sensitivity and specificity of GVTD to motion. To facilitate comparisons between the accelerometer and GVTD, the angular rotation was calculated based on the final head orientation time-course (§2.5, Fig. 3I).

We evaluated the efficacy of GVTD and angular rotation for motion detection through different scenarios. First, we compared these two motion indices for a gross and a small artifact and found that GVTD shows a higher amplitude spike than the angular rotation in the case of small artifacts such as eyebrow motion (Fig. 4A, B). To quantify these comparisons, we first calculated the Pearson correlation between GVTD and angular rotation (**Φ**) for all runs containing instructed motion. The correlations were averaged over the six subjects that had concurrent HD-DOT and motion sensor data for all runs in the session (Fig. 4C). These correlations were greatest in cases of head rotations (r = 0.86 ± 0.06 for roll and pitch) and lowest for eyebrow motion (r = 0.46 ± 0.2). This difference most likely reflects the transducer characteristics of the motion sensor and the fact that it is not sensitive to the small muscle movements when attached to the top of the HD-DOT cap.

**Figure 4.**
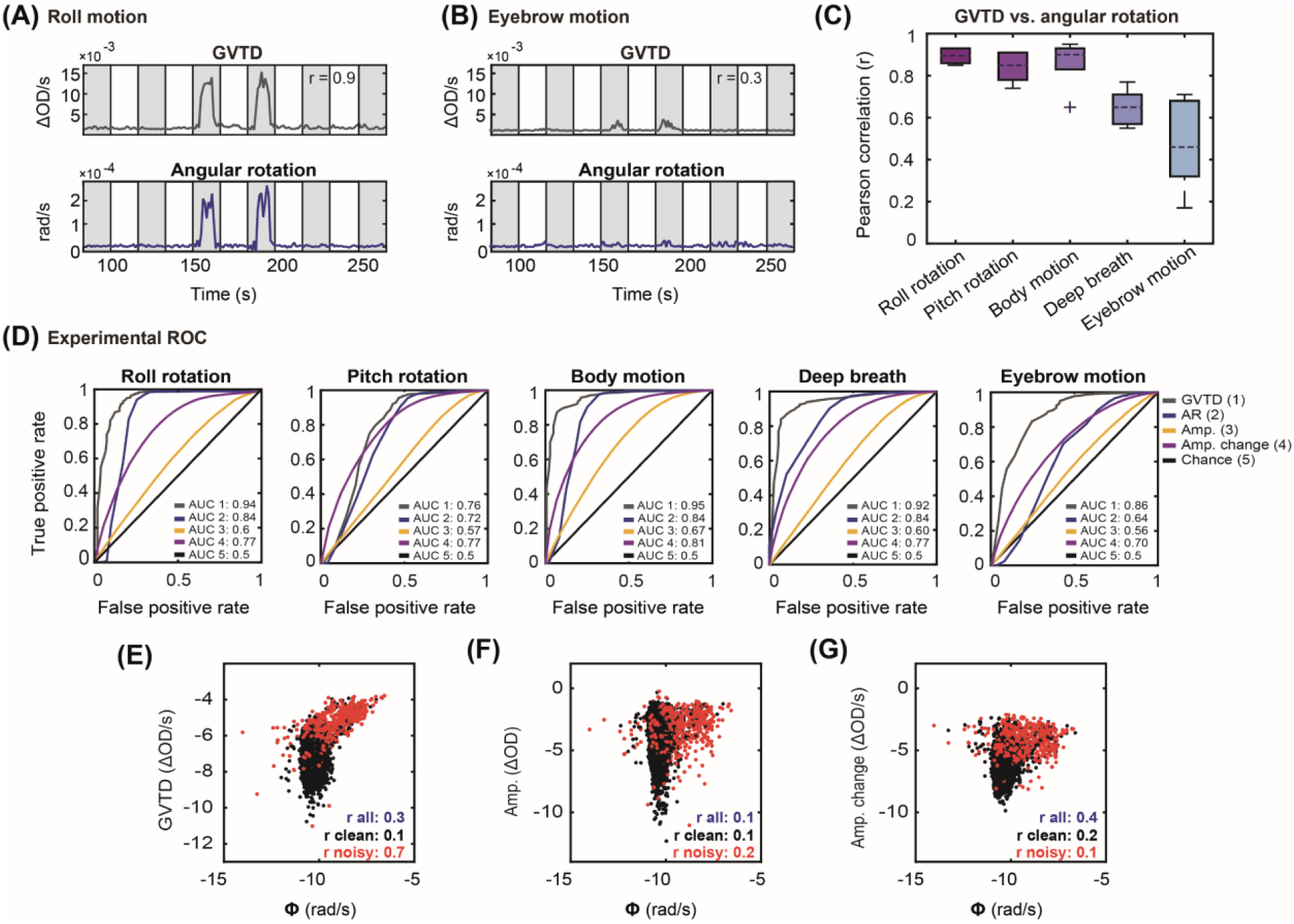
GVTD and angular rotation for an example HW run containing instructed (A) roll and (B) eyebrow motion artifacts. Gray shaded regions indicate auditory stimulation. (C) Pearson correlation between GVTD and angular rotation averaged over the six subjects with instructed motion runs. Note high Pearson correlation of GVTD with roll, pitch, and body motion. (D) Experimental ROC plots for GVTD and angular rotation and the mean of the ROCs for signal amplitudes (Amp.) and windowed amplitude changes (Amp. change) for five types of instructed motion. Log-log scatter plots of (E) all 850 nm nn1 signal amplitudes, (F) all 850 nm nn1 windowed amplitude changes, and (G) GVTD vs. angular rotation for all runs with instructed motion. The correlation between the GVTD and the motion sensor is higher than both amplitudes and the windowed amplitude changes. The cutoff between black and red dots is based on the instructed motion time-points.

To evaluate the sensitivity of the GVTD to motion, we leveraged the ground truth built into our instructed motion paradigm. Experimental receiver operator characteristic (ROC) curves for GVTD and angular rotation were created for a binary classification of clean and noisy time-points by sweeping the detection threshold (Fig. 4D). We defined ground truth for motion as the time-points during which the subjects performed instructed movements. We also plotted these ROC curves for two common temporal motion detection methods in fNIRS, i.e., absolute single-channel signal amplitudes and windowed amplitude changes for all motion types and all 850 nm nn1 measurements (Fig. S3) and compared the mean of these ROC curves against GVTD and angular rotation (Fig. 4D). In all motion types, GVTD showed better or similar performance (AUC) compared to angular rotation, absolute signal amplitude, and windowed amplitude change (Table 3).

**Table 2:** Percent of the noisy measurements (nn1 through nn4) across five different instructed motion artifacts in dataset 2. Noisy measurements were empirically defined as ones having a temporal standard deviation of 7.5% or greater.

| Type of motion | Roll | Pitch | Deep breaths | Body motion | Eyebrow motion |
| --- | --- | --- | --- | --- | --- |
| % Noisy measurements | $49 \pm 9\%$ | $47 \pm 4\%$ | $41 \pm 5\%$ | $51 \pm 8\%$ | $39 \pm 4\%$ |

**Table 3:** The area under the curve (AUC) of the experimental receiver operator characteristic (ROC) of GVTD and angular rotation (based on the motion sensor outputs), the mean of the ROC of the absolute signal amplitude and windowed amplitude changes based on the instructed motion as ground truth in dataset 2.

| Motion index | GVTd | Angular rotation | Signal amplitude | Windowed amplitude change |
| --- | --- | --- | --- | --- |
| AUC | $0.88 \pm 0.07$ | $0.77 \pm 0.08$ | $0.6 \pm 0.04$ | $0.76 \pm 0.04$ |

We used the instructed motion protocol to examine the relation between GVTD and angular rotation for all runs with instructed motion (Fig. 4E). Low vs. high motion time-points (black vs. red in Fig. 4E) were determined based on the ground truth of the instructed motion protocol (high motion as defined as time-points when the subject performed instructed motion). When the motion was low (black dots) GVTD and angular rotation were not correlated (r = 0.05 ± 0.05), but when the motion was high (red dots), GVTD and angular rotation were highly correlated (r = 0.8 ± 0.1). The same log-log scatter plots for absolute signal amplitudes (Fig. 4F) and the windowed amplitude (Fig. 4G), show much lower correlations with the angular rotation (0.2 and 0.1, respectively) compared to GVTD (0.7).

In summary, these results show that GVTD can be used as an alternative or in conjunction with motion sensors in detecting noisy time-points of data.

### 3.3 Motion detection strategy using GVTD

To censor data using the GVTD time-course, we developed an outlier detection strategy that separates good data from motion artifacts.

We assume that the detected signal, *y*(*t*), is a linear combination of the true physiological signal, *S*(*t*), and noise, *ϵ*(*t*):

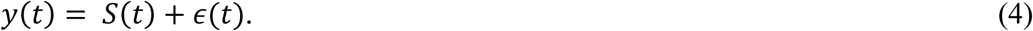

We followed the fMRI approaches for DVARS and FD and developed a data distribution driven strategy for finding motion criterion. In fMRI, *S*(*t*) is approximately normally distributed [56]. Accordingly, the DVARS distribution is right-skewed [57]. Therefore, we investigated the skew of the GVTD distribution as a potential index of head motion artifact in HD-DOT. We evaluated the GVTD distribution for HD-DOT data from a still Styrofoam phantom, a low motion trial, and a high motion trial. The phantom GVTD histogram peaked at a relatively small value (mode = 4 × 10^−5^) and exhibited a small rightward skew (Fig. 5A). In the low motion human data, GVTD values had a higher mode and proportionately smaller skew (Fig. 5B). In data with instructed motion (high motion), the GVTD distribution is strongly skewed to the right (Fig. 5C). These results suggest that the skew provides a basis for censoring HD-DOT data.

**Figure 5.**
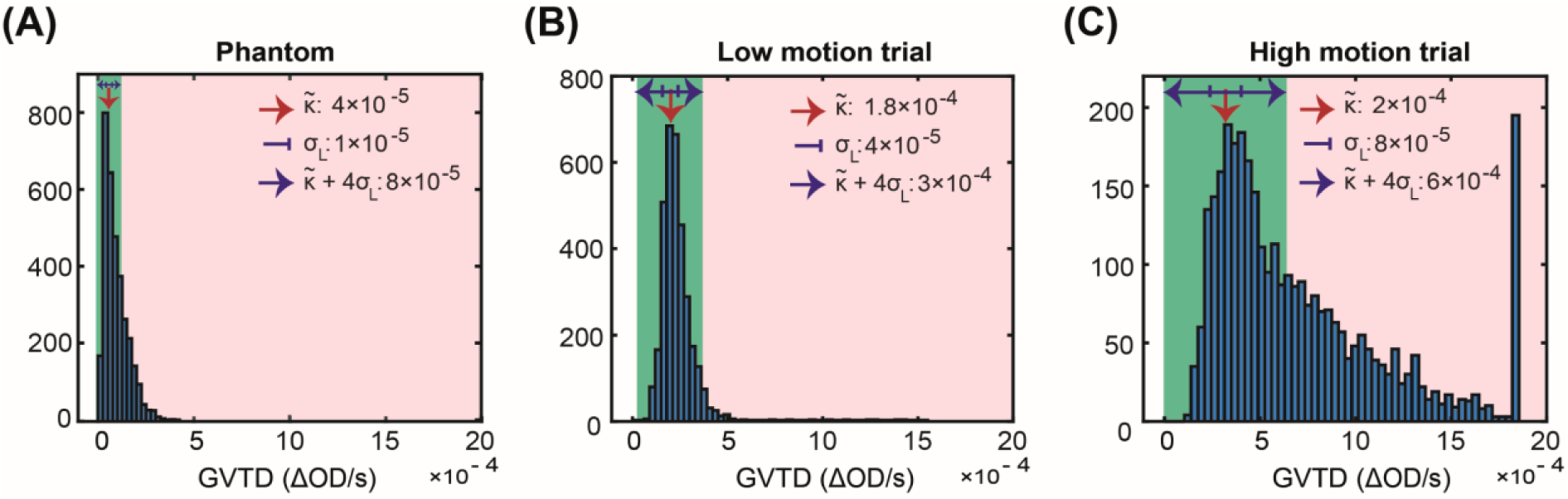
Histograms of the GVTD values for (A) a Styrofoam phantom, (B) a low motion run, and (C) a high motion run in one subject. Note the constant x-axis limits; values above that limit fall into the last bin. Note mode (red arrow), left standard deviation, *σ*_*L*_, and noise threshold computed according to Eq. 5. Pink and green shading indicate GVTD values that do or do not exceed the noise threshold.

Thus, we defined a noise threshold (*g*_*thresh*_) based on the GVTD distribution mode 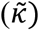 plus a constant (*c*) times the standard deviation computed on the left (low) side of the mode (*σ*_*L*_). The right tail of the GVTD distribution corresponds to motion artifacts (Eq. 5). Thus,

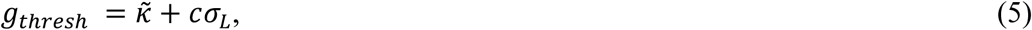

where 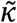 is the histogram mode and *σ*_*L*_ is computed as 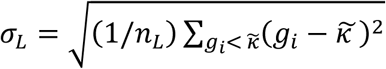, where *n*_*L*_ is the number of GVTD time-points less than 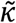. The value of *c* controls the trade-off between the exclusion of artifact vs. data loss.

### 3.4 Determining the best stage for performing GVTD-based motion detection and censoring

GVTD is a generic measure that can be applied to any data in the form of channels (or voxels) by time. Therefore, we needed to determine where in the processing pipeline, GVTD should be performed. We evaluated four potential locations (green boxes in Fig. S1). To evaluate GVTD’s ability to separate noise from the signal, we defined an artifact-to-background ratio (ABR) as the mean of the GVTD values above a noise threshold over the mean of the GVTD values below the threshold. Specifically, GVTD was calculated for; **a)** SD-pair log-mean optical densities *(“after log-mean”*; unit = optical density change per second (Δ*OD*/*s*)), **b)** after temporal filtering before superficial signal regression (SSR) (“*after filtering no SSR*”; unit = Δ*OD*/*s*), **c)** after both temporal filtering and SSR (“*after filtering with SSR*”; unit = Δ*OD*/*s*), and **d)** on reconstructed image voxels (“*after reconstruction*”; unit = molar HbO_2_/s). These results were compared based on their ABR means on two different datasets with natural motion to determine the most effective GVTD strategy. GVTD time-courses, GVTD histograms, and their associated gray plots calculated at these four stages for resting state data collected with two HD-DOT systems (example of a run from the adult HD-DOT data in Fig. 6A-C). The ABR index (Eq. 3) was calculated using the motion threshold defined as 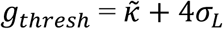 (Eq. 5). Results showed that ABR was consistently highest after both filtering and superficial signal regression but before image reconstruction in both datasets 4 and 5 (Fig. 6D).

**Figure 6.**
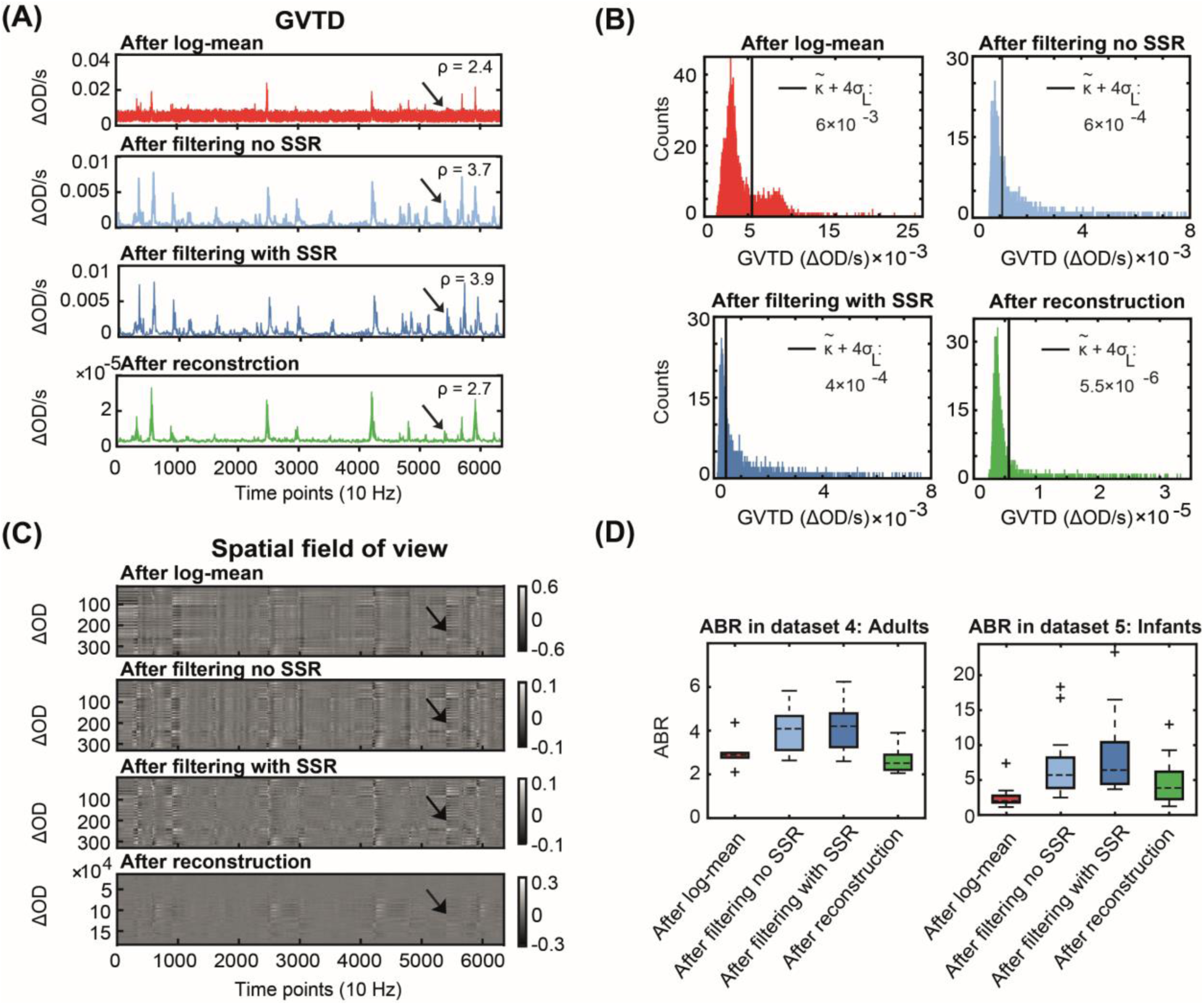
Determination of optimal GVTD stage in the processing pipeline based on the artifact-to-background ratio (ABR; Eq. 3) of the 850 nm first nearest neighbor measurements. (A) GVTD computed after log-mean, after filtering and without SSR, after filtering and with SSR, and after reconstruction. (B) Histograms of the GVTD values for the four time-courses; black lines indicate the noise threshold 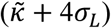. (C) Four gray plots associated with the four GVTD time-courses shown in (A). Black arrows indicate a small motion artifact. Note the greatest contrast between the motion artifact and the baseline after filtering (third time-course). (D) ABR values calculated for all four processing stages for all subjects in datasets 4 and 5. GVTD after filtering (light and dark blue) was maximal in all cases and the highest after filtering with SSR (dark blue).

### 3.5 Indexing data quality with GVTD in task HD-DOT data by comparison against fMRI

Dataset 2 was used to evaluate the ability of GVTD to index the HD-DOT data quality. HD-DOT responses to hearing words were compared to the group-mean fMRI response to the same task, which was independently acquired in a separate experiment and treated as a “gold standard”. We rank-ordered ordinary HD-DOT trials for each subject according to their mean GVTD value; for each subject, the ten lowest and ten highest GVTD ordinary trials were defined as “low motion” and “medium motion”. The instructed motion trials were defined as “high motion”. Responses were extracted from a fixed ROI defined as P < 0.05 in the fMRI dataset (Fig. 8A, 3^rd^ column map), expressed as percent signal change. The Pearson correlation between the HD-DOT and fMRI time-courses were computed for each of the three HD-DOT conditions (Fig. 7B-D). This correlation progressively decreased from 0.97 for low motion to 0.86 for medium motion, to 0.78 for instructed motion (Fig. 7E). Medium motion responses (Fig. 7C) were comparable to fMRI, but with a smaller peak value and higher mean squared error (0.08). Trials that GVTD identified as low motion (Fig. 7B) generated the cleanest maps with the lowest mean squared error (0.06). Accordingly, the GLM-derived beta-values were greater in the low as compared to high motion trials in most subjects (Fig. 7G).

**Figure 7.**
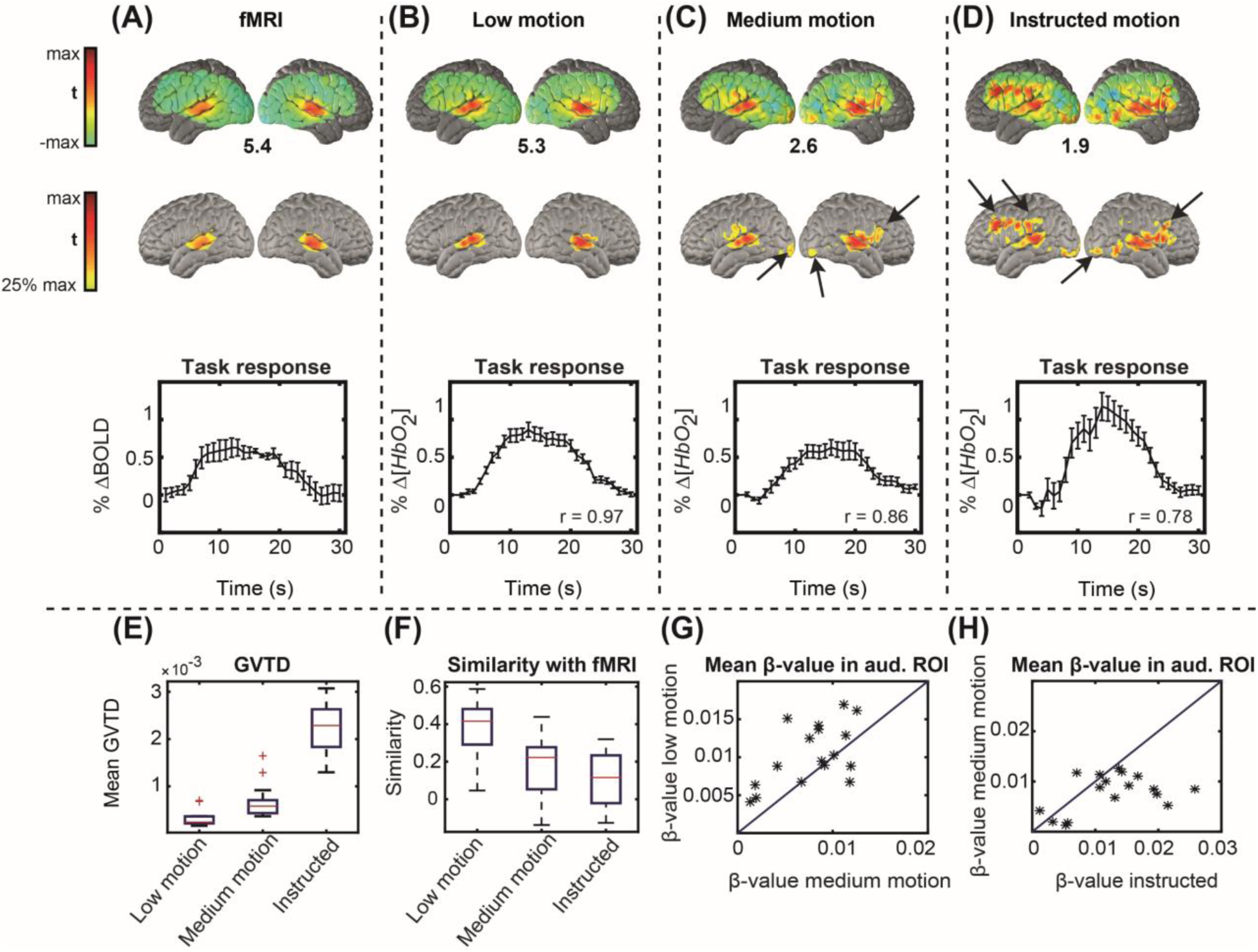
HW task t-statistic evoked responses. Voxel-wise maps are shown in the first and second rows; the percent signal changes are averaged over a region of interest (ROI) and shown in the 3^rd^ row. Error bars on the time-courses indicate standard error of the mean across sessions. (A) Reference dataset. (B) Low motion data. (C) Medium motion data. (D) Instructed motion data. Black arrows indicate false positive responses, designated since they occur outside auditory ROI defined based on the reference fMRI dataset. (E) Mean GVTD values across all trials in low motion, medium motion, and instructed motion data. (F) Mean similarity of the maps in each condition with the reference dataset, similarity defined as the voxel-voxel Pearson correlation. (G) Scatter plot of responses in low vs. medium motion ordinary trials; GVTD indexed stronger responses in low motion trials in 15 of 17 sessions. (H) Scatter plot of medium motion vs. instructed motion trials; note the higher spurious response magnitudes for the instructed motion.

**Figure 8:**
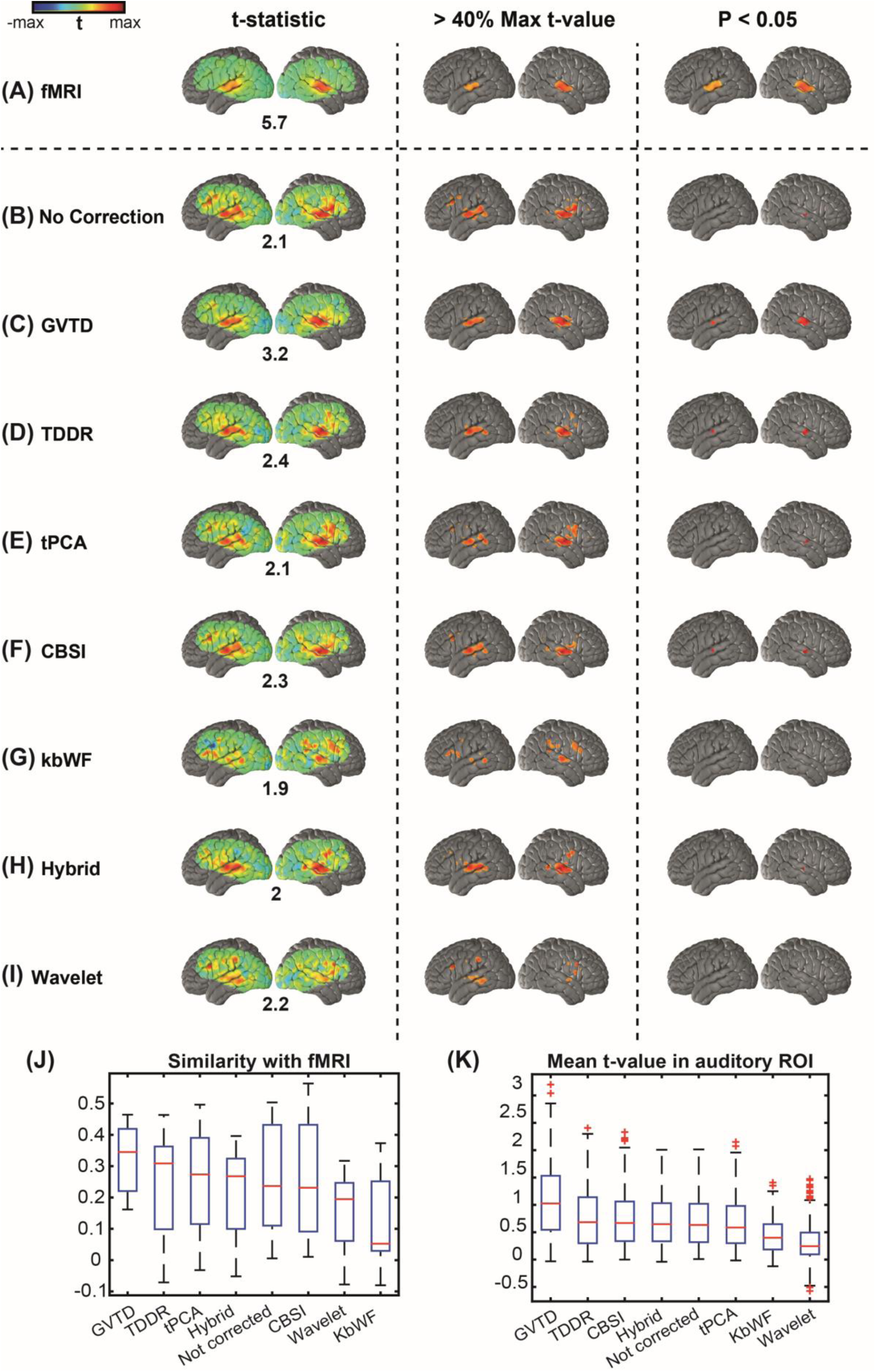
Comparison between different motion removal methods on HW task HD-DOT data in dataset 3 based on the gold standard fMRI map from dataset 1. Three columns represent the same t-statistic map for each row with: 1. No threshold, 2. thresholded at 40% of the maximum t-value of each method (mapped as an alternative visualization), and 3. Thresholded based on the P < 0.05 statistical significance. (A) fMRI maps based on reference dataset 1. HW map for dataset 3 with (B) no motion correction. (C) GVTD-based motion censoring, (D) TDDR (E) tPCA (F) CBSI (G) kbWF (H) hybrid, and (I) wavelet motion correction methods. (J) The similarity between the non-thresholded t-statistic maps is calculated based on the voxel-voxel Pearson correlation with fMRI t-statistic map. (K) The mean t-value is calculated in the auditory ROI based on the fMRI HW map thresholded at P < 0.05 shown in panel A column 3.

A cautionary point regarding GLM-derived beta values is raised by the instructed motion trials, which generated the highest mean squared error (0.12) as well as the greatest BOLD response modulations, hence, the greatest GLM-derived beta values (Fig. 7H). These response time-courses were the least similar to those obtained by fMRI (Fig. 7F) and were accompanied by voxel-wise activations outside of the auditory cortex. Thus, the apparently strong HD-DOT responses in the instructed motion condition are attributable to motion artifact, as detected by GVTD (Fig. 7E). We conclude that the results shown in Fig. 7 demonstrates that GVTD effectively indexes HD-DOT data quality.

Additional results derived from the HW response analysis show a progressively lower similarity of the HW responses for fMRI results in association with greater GVTD values (Fig. 8 E, F). The relationship between low motion and medium motion data within each session shows that responses are systematically greater in low motion as opposed to medium motion (true in 15 out of 17 sessions). The responses are comparably compromised by spontaneous motion in medium motion trials (as indexed by greater GVTD) and spuriously higher in instructed motion trials with the highest GVTD scores (Fig.7 G, H).

### 3.6 Comparison between motion removal methods applied to HW task HD-DOT data

To compare the performance of different motion removal methods on HD-DOT data, we used dataset 3, acquired in older subjects (n = 13; 42 ± 19.75 years old) performing the hearing words task (no instructed motion). Dataset 3 included a wide range of motion contamination levels. The details of the various motion removal methods used in this analysis are explained in §2.3. Responses were evaluated in terms of statistical significance at the voxel and ROI levels as well as time-course similarity with fMRI.

Without motion removal, the group-level t-statistic map contained several spurious activations that are not present in the fMRI results (Fig. 8A, B). Moreover, the expected superior temporal cortex response did not achieve statistical significance at P < 0.05. In this analysis, the GVTD threshold was computed as 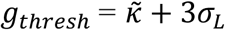 (Eq. 5). Exemplary low and motion and high motion blocks are illustrated in Supplementary Fig. S4. This threshold excluded all blocks in 6 subjects, leaving 7 subjects contributing to the final result illustrated in Fig. 8C. Results obtained with TDDR, tPCA, CBSI, kbWF, hybrid (Spline + Savitzky Golay), and wavelet filtering are illustrated in Fig. 8D-I. GVTD censoring, TDDR, and CBSI methods recovered bilateral superior temporal cortex activations in thresholded t-statistic maps (P < 0.05). tPCA and hybrid methods also recovered a unilateral right hemisphere activation. However, no statistically significant (P < 0.05) responses were obtained with the other methods (wavelet and kbWF).

We quantified the performance of the results shown in Fig. 8B-I using two metrics: 1. Similarity score, defined as the voxel-wise Pearson correlation between the non-thresholded maps and the fMRI gold standard map, and 2. Mean t-value in the auditory ROI defined P < 0.05 in the fMRI t-map (Fig. 8A, 3^rd^ column). The spatial similarity to fMRI was greatest for the GVTD-censored map, followed by TDDR, tPCA, hybrid, not-corrected, CBSI, wavelet, and kbWF maps (Fig. 8J). The mean ROI t-value was greatest for the GVTD-censored maps, followed by TDDR, CBSI, hybrid, not corrected, tPCA, kbWF, and wavelet corrections (Fig. 8K). As noted above in §3.5, artifacts can spuriously increase apparent response magnitudes, hence, GLM-derived t-values. This observation underscores the value of comparing HD-DOT results to those of fMRI.

### 3.7 Comparison between motion removal methods applied to resting state HD-DOT data

We compared the performance of different HD-DOT motion removal methods in application to resting state HD-DOT data using dataset 4 (n = 8 adults, 30.25 ± 11.18 years old). Seed-based functional connectivity (FC) was computed using the 5 seed ROIs (§2.9.4, Fig. 9 top row). In parallel with §3.6, we quantified the performance of each correction method using two metrics: 1. similarity score, defined as the spatial similarity between the HD-DOT and fMRI FC maps; and 2. Mean FC (Fisher z-transformed correlation) in functionally connected ROIs identified in the fMRI data. The spatial similarity was computed as the Fisher z-transformed Pearson spatial correlation between non-thresholded maps, evaluated over the HD-DOT FOV (white area illustrated in the top row of Fig. 9). Mean FC was evaluated in the colored ROIs illustrated in Fig. 9A. Thus, this measure reflected simple homotopic FC in primary cortical areas as well as ipsilateral FC in the higher-order networks (DAN and FPC). The GVTD threshold was computed as 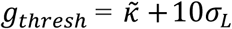 (Eq. 5). This lenient threshold minimized data loss. On the basis of preliminary testing, GVTD censoring was extended to retain only epochs of duration at least 30 seconds.

**Figure 9:**
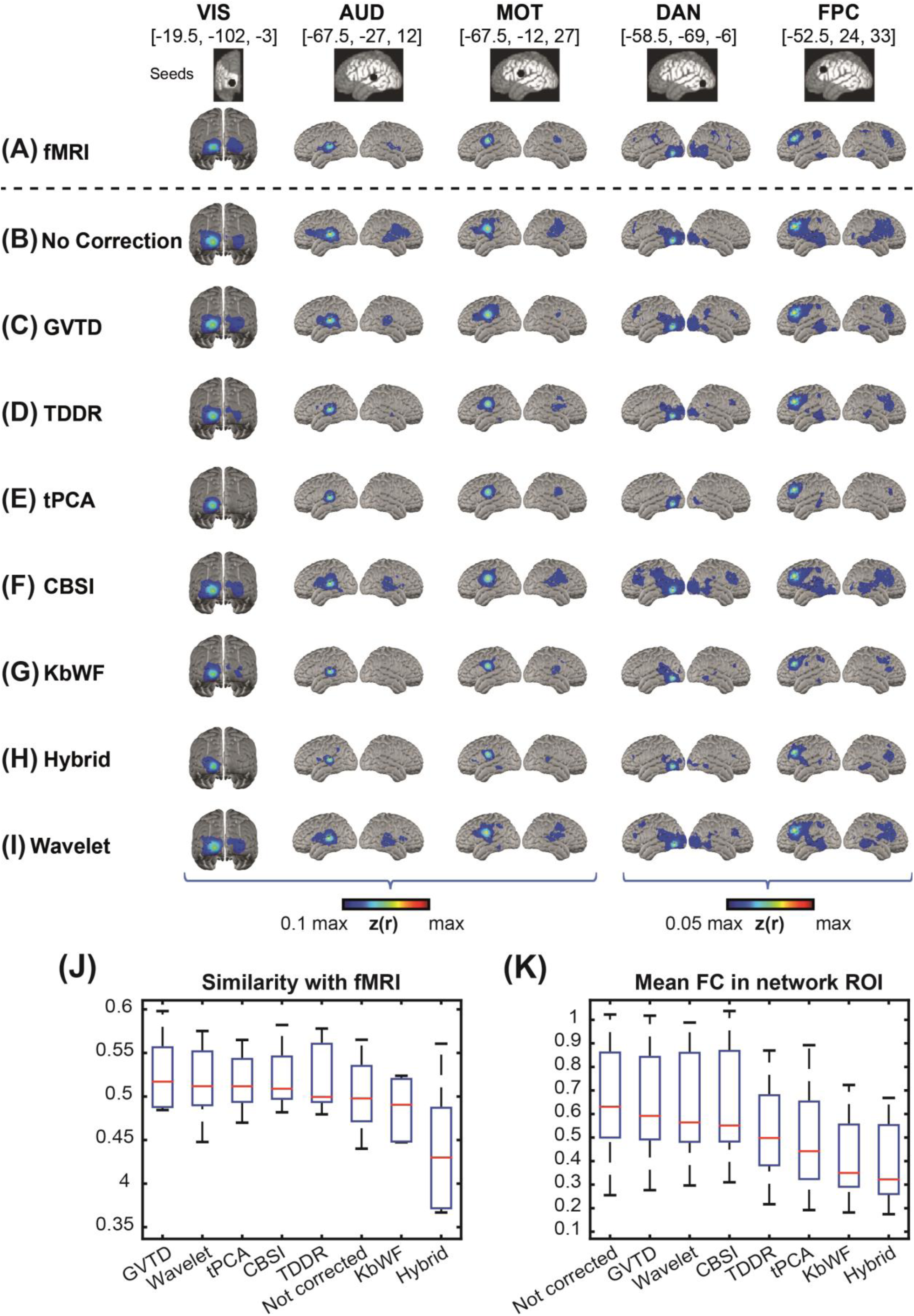
Comparison between different motion removal methods on resting state HD-DOT data in dataset 4 and gold standard fMRI map. Five columns represent the seed maps for visual (VIS), auditory (AUD), somatomotor (MOT), dorsal attention (DAN), and frontoparietal (FPC) networks. The first 3 seed maps are thresholded at 10% of the maximum z(r) value of each category and the last 2 higher order networks (DAN and FPC) are thresholded at 0.5% of the maximum z(r) value of the group maps. (A) fMRI maps based on reference dataset 1. HD-DOT maps for dataset 4 with (B) no motion correction. (C) GVTD-based motion censoring, (D) TDDR (E) tPCA (F) CBSI (G) kbWF (H) hybrid, and (I) wavelet motion correction methods. Spatial similarity (J) was computed as the Fisher z-transformed spatial correlation between the HD-DOT and fMRI FC maps, evaluated over the HD-DOT FOV (white area illustrated in the top row). ROI-based FC (K) was evaluated as the mean Fisher z-transformed correlation with the seed in the colored regions shown panel A. These regions were determined by thresholding the group-level fMRI FC maps at 10% (for lower level networks (VIS, AUD, and MOT) and 5% for higher level networks (DAN and FPC) of Fisher’s z-score.

The results obtained by the various correction methods are shown in Fig. 9B-I. The most extensive HD-DOT FC maps were obtained in uncorrected data (Fig. 9B). However, these maps were not spatially most similar to the fMRI gold standard dataset. Rather, GVTD censoring (Fig. 9C) yielded HD-DOT FC maps most similar to fMRI (Fig. 9J). Of all censoring methods, GVTD yielded the greatest FC in the evaluation ROIs, followed by wavelet, CBSI, TDDR, tPCA, kbWF, and hybrid corrections (Fig. 9K). As in the HW task responses, strong FC in the evaluation of network ROIs does not necessarily indicate good data quality, especially when accompanied by spurious effects outside of the network identified on the basis of fMRI (e.g., as seen in the no correction, wavelet, and CBSI maps). On the other hand, some methods may overcorrect, leading to falsely weak correlations (TDDR, tPCA, kbWF, and hybrid methods).

## 4 Discussion

### 4.1 A general summary of the novel strategies and findings

We developed a novel motion detection method suitable for high-density optical imaging arrays, inspired by the DVARS in fMRI [43]. Specifically, we defined the global measure of variance in the temporal derivative across measurement channels (GVTD) and developed a method for denoising structured artifacts in HD-DOT. We found that GVTD successfully indexes motion artifacts in HD-DOT and has higher sensitivity and specificity (evaluated using AUC of the ROC curve against the ground truth of instructed motion) for motion detection compared to an accelerometer motion sensor and to single-channel motion detection methods commonly used in fNIRS (absolute signal amplitudes and windowed amplitude changes).

While there are a number of papers evaluating motion removal methods for standard fNIRS [34-39, 46, 58-61], the literature on motion removal strategies for HD-DOT is limited. Previous studies lack some combination of HD-DOT datasets and comparisons to gold standard data (fMRI) for image quality validations and most are restricted to single-channel motion detection. In this paper, we introduce a novel approach for evaluating the efficacy of motion removal methods in HD-DOT by comparison against matched fMRI datasets.

We show that the mean GVTD score is correlated with the similarity of the HD-DOT task images to those of fMRI. Thus, the mean GVTD score can be used to classify datasets as either clean or noisy (Fig. 7). We also show that applying GVTD censoring to both task and resting state HD-DOT datasets outperforms other fNIRS-based motion correction methods and makes HD-DOT maps more similar to those of fMRI. Together, HD-DOT imaging arrays and anatomical atlasing combined with GVTD motion censoring, all aid in making HD-DOT data more comparable to fMRI and furthers the use of HD-DOT as a surrogate to fMRI.

### 4.2 Optimizing the implementation of GVTD in the HD-DOT processing pipeline

We optimized the use of GVTD motion detection in HD-DOT by testing it at different steps of the processing pipeline using the artifact-to-background ratio (ABR). In fMRI, DVARS has only been evaluated before and after filtering [55]. In contrast, in HD-DOT, we can consider GVTD in either measurement space or image space (after image reconstruction). Our results show that the ABR was highest in measurement space prior to image reconstruction and after filtering the high-frequency content of the data. It was also statistically better when performed after SSR, a common fNIRS and DOT processing step (in datasets 4 and 5). Therefore, based on our ABR analysis, we recommend performing GVTD after filtering the measurements, but prior to image reconstruction.

An important decision with GVTD is to determine the censoring threshold. Since the baseline GVTD value was different across people, similar to findings with DVARS in fMRI [50], we evaluated a noise detection strategy based on the GVTD distribution (histogram) specific to each subject. The differences in the baseline GVTD distribution is possibly due to variable physiological signal levels as well as respiratory patterns, heart rate, facial muscle activity, restlessness, tremor, etc. [21, 55]. Therefore, we developed an outlier detection strategy individualized for each subject’s data that semi-automates the noise threshold determination and takes into account subject differences. Specifically, we set the threshold using the GVTD distribution mode 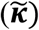 plus a constant (***c***) determined based on the left side (lower side) of the mode of the GVTD distribution. For practical implementation, we recommend that the threshold be greater than the standard deviation of the baseline signal.

### 4.3 Evaluation and validation of denoising through comparisons to fMRI

Most fNIRS studies measure the efficiency of motion removal techniques based on the recovery of a synthetic HRF [31-33, 36], or, in the case of real data, based on the variance across subjects or datasets [33]. However, since HD-DOT is focused on creating images comparable to fMRI, throughout this paper, we have used an fMRI dataset with the same task and resting state paradigm used in our HD-DOT datasets as a gold standard for evaluating the efficacy of different motion removal methods. The comparisons were based on the voxel-voxel Pearson correlation of the spatial HD-DOT maps and the fMRI maps. We find that, for both task and resting state functional connectivity, comparisons to fMRI enables identifying false negatives, false positives, and localization errors (Figs. 8, 9), all of which would be difficult to determine without a target image. In vivo imaging enables a much stronger evaluation than in silico simulations; fMRI data contains real image features, including the spatial extent, signal magnitude, distribution of spatial frequencies, and time-courses.

Using the fMRI comparisons, we ranked ordered several motion removal methods in both task and resting state data. The general pattern observed was that motion censoring using GVTD worked best, with near contenders being CBSI, TDDR, and following those, targeted PCA in both task and rest data. TDDR and tPCA both suppressed the mean t-value in the auditory ROI and FC in the evaluation ROIs, which may indicate overcorrection, i.e., removal of the true signal. Wavelet filtering ranked second after GVTD censoring in resting state data both in terms of similarity with fMRI and mean FC in the resting state networks. This is in distinction to its lower performance in the task data.

### 4.4 On different performance of motion correction methods in fNIRS literature

A striking aspect of the fNIRS literature is the variable performance of motion correction methods across different studies [31-33, 35, 36, 39]. One possible reason for the variability between the studies could be the different levels of motion present in each study. This variability has also been evaluated in a recent fNIRS study [33]. To address this topic, we performed a supplementary analysis of the low motion, medium motion, and high motion HW task data in dataset 2 (Fig. 7). We evaluated the performance of different motion correction methods on different levels of motion artifacts in these three categories (Figs. S5, S6).

This analysis shows that, in the low motion group, all methods can preserve bilateral auditory cortex HW responses. In the medium and instructed motion groups, GVTD, TDDR, CBSI, and tPCA again outperformed other methods by recovering either a unilateral or bilateral HW activation with no obvious false positives in the P < 0.05 thresholded maps (Fig. S7). Note that, in the high motion data (instructed motion group), none of the motion correction techniques fully recovered bilateral auditory responses (present in fMRI). However, GVTD was able to distinguish between clean vs. motion-corrupted data (Fig. 7). We hypothesize that GVTD can provide a means of rank-ordering data based on quantitative motion estimation (as suggested in Fig. 7), something that is normally done subjectively prior to applying motion correction methods. Thus, GVTD may be useful also in denoising sparse fNIRS data. This notion could be tested by evaluating the efficacy of GVTD in sparse fNIRS arrays or by subsampling the HD-DOT imaging array.

GVTD focuses on motion detection, followed by simple censoring. GVTD could be used as an alternative to either absolute signal amplitudes or windowed amplitude changes included in the Homer2 code package [62]. Further, GVTD could be used in conjunction with motion correction methods such as spline interpolation (MARA) [34], Kalman filtering [58, 63], PCA [58], tPCA [46], Hybrid methods [36], or any method that depends on motion detection in the temporal domain. However, we note that, in the results presented here, GVTD-based censoring alone provided better image quality than any of the alternative motion correction procedures.

### 4.5 Strengths and limitations of the GVTD-based motion censoring

When tested in HD-DOT, the most promising results were obtained using GVTD-based motion censoring. A likely reason for GVTD efficacy is that it leverages the effect of small artifacts across many measurements. The simplicity of GVTD censoring guarantees that the signal is neither over-smoothed nor overcorrected.

As described here, GVTD is used as a binary classifier to censor the time-points marked as noisy. However, it also could work with a non-binary weight associated with the time-points based on their GVTD value to soften the impact of threshold choice. For example, time points with GVTD values closer to the GVTD distribution mode could be assigned higher weights than ones further from the mode [64].

Another important challenge in scrubbing data is the tradeoff between losing signal vs. removing noise [65]. For motion criterion, one can ensure that sufficient data remains after censoring by tuning *c* (Eq. 5). Another approach would be to use GVTD to determine the useable data yielded from a run and then adjust the data collection to either collect more data within the session or add sessions to the study. These active data quality approaches are currently being pioneered in fMRI with runtime assessment of motion [66-68].

### 4.6 Summarizing the consensus regarding the top-performing denoising strategies in the fNIRS literature

Among the fNIRS-based methods that worked best for HD-DOT, besides GVTD, CBSI performed well in both task and resting state data. CBSI does not require tuning of parameters but has been less recommended in the literature [35, 36] as it relies on the assumption of a negative correlation between HbO and HbR. Therefore, it is limited to populations in which a normal correlation between HbO and HbR can be assumed [37].

The TDDR method performed well in the task HD-DOT data and fairly well in the resting state analysis. TDDR, like CBSI, does not require tuning of parameters. However, one disadvantage of TDDR is that it relies on the derivative of single measurements and, thus, is less sensitive to small motion artifacts such as eyebrow motion. Moreover, TDDR only performs an efficient motion correction on the low-frequency content of the data, because the higher frequencies inflate the variance of the temporal derivative distribution and create bias in the distribution of estimates [35]. However, we showed that the noise content is still present in the data after band-pass filtering (see post-filtering gray plots in Fig. 6C showing residual artifact during motion).

Targeted PCA also yielded HD-DOT maps similar to those in fMRI but with decreased response magnitudes in both task and resting state data. tPCA removes a fixed proportion of variance through the removal of the largest principal component; hence, as observed here, is prone to overcorrection [35, 46].

Wavelet filtering, despite a poor performance in task data, showed good performance in resting state HD-DOT data. However, this method is computationally expensive. On average, for both HW and rest HD-DOT runs, wavelet filtering ran ten times slower than other motion correction or censoring methods. The kbWF method, while faster than the full wavelet approach, did not perform well in either task or rest HD-DOT data.

## 5 Conclusion

We developed GVTD, a novel motion detection metric and optimized its use in the HD-DOT pre-processing pipeline. GVTD can be used alone or in combination with other motion correction methods to increase the quality of data obtained with multi-channel optical imaging systems. We evaluated GVTD using several independent HD-DOT datasets, including an instructed motion protocol, accelerometer motion measures, and a matched fMRI dataset serving as ground truth. Although GVTD-based censoring removes data, the obtained HD-DOT maps were most similar to those of fMRI and it outperformed alternative motion correction methods previously described in the fNIRS literature.

## Acknowledgments

The authors would like to acknowledge the following funding resources: NIH - R21NS098020 (JPC and TH), R21DC016086 (JPC), U01 EB027005 (JPC and ATE), U54HD087011 (JPC), R01 NS090874 (JPC), K01MH103594 (ATE), R21MH109775 (ATE), KL2 TR000450-7 (BJP), 1P30NS098577 (AZS and JPC), 1P01NS080675-01A1 (AZS, TH, and J.P.C), K02 NS089852 (CDS), UL1 TR000448 (CDS and JPC), and Mallinckrodt Institute of Radiology (CDS and JPC). AS wants to acknowledge helpful discussions with Jonathan E. Peelle, Broc A. Burke, Zachary E. Markow, Andrew K. Fishell, and Kalyan Tripathy.

## 6 Data availability

The data that support the findings of this study are available from the corresponding author upon reasonable request.

## 7 Conflict of interest

*The authors declare that the research was conducted in the absence of any commercial or financial relationships that could be construed as a potential conflict of interest*.

## Supporting materials

The following supporting materials can be found online in the supporting information tab for this article.

Supplementary Figures

Supplementary Tables

