## Supplementary Figures for "Global Motion Detection and Censoring in High-Density Diffuse Optical Tomography"

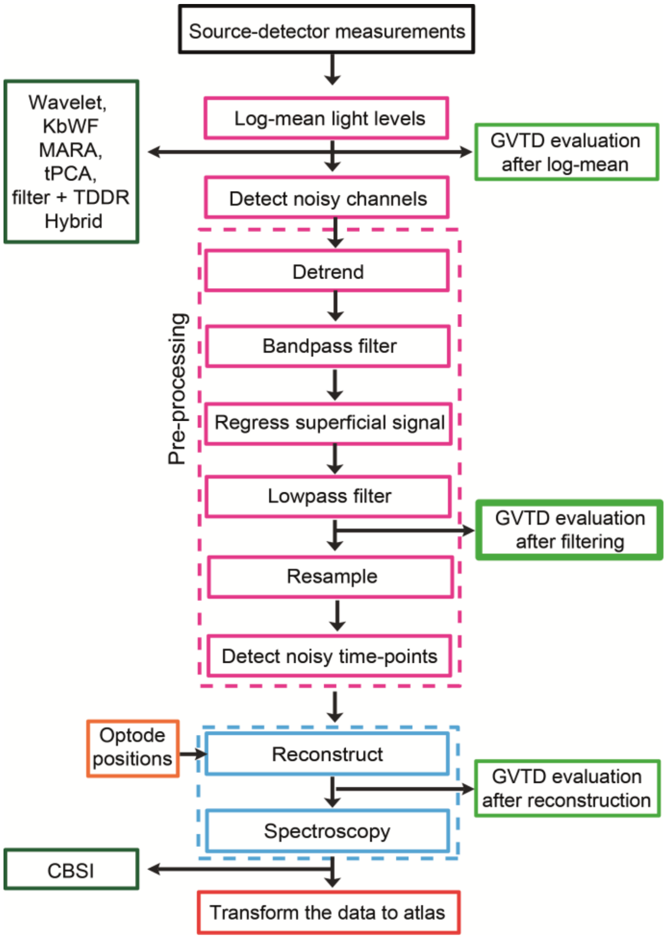


**Figure S1.** Processing of HD-DOT data. Dashed pink and blue boxes show the pre- and post-processing steps, respectively. GVTD was evaluated (section 3.4) at three stages in the pipeline indicated by green boxes. The best separation between baseline signal and motion artifact was obtained "after filtering" (thicker outlined green box).

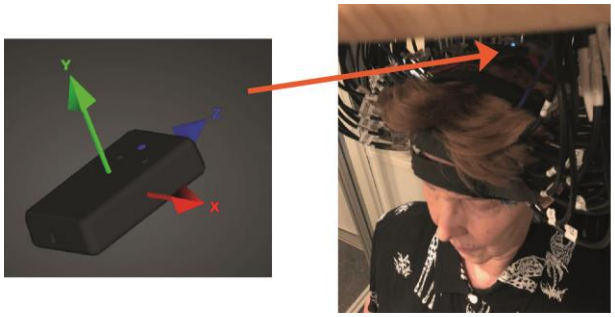

**Figure S2.** The motion sensor (3-space™ USB/RS232; Yost Labs) was attached to the top strap of the HD-DOT device for concurrent recording of the head motion and optical data in dataset 2.

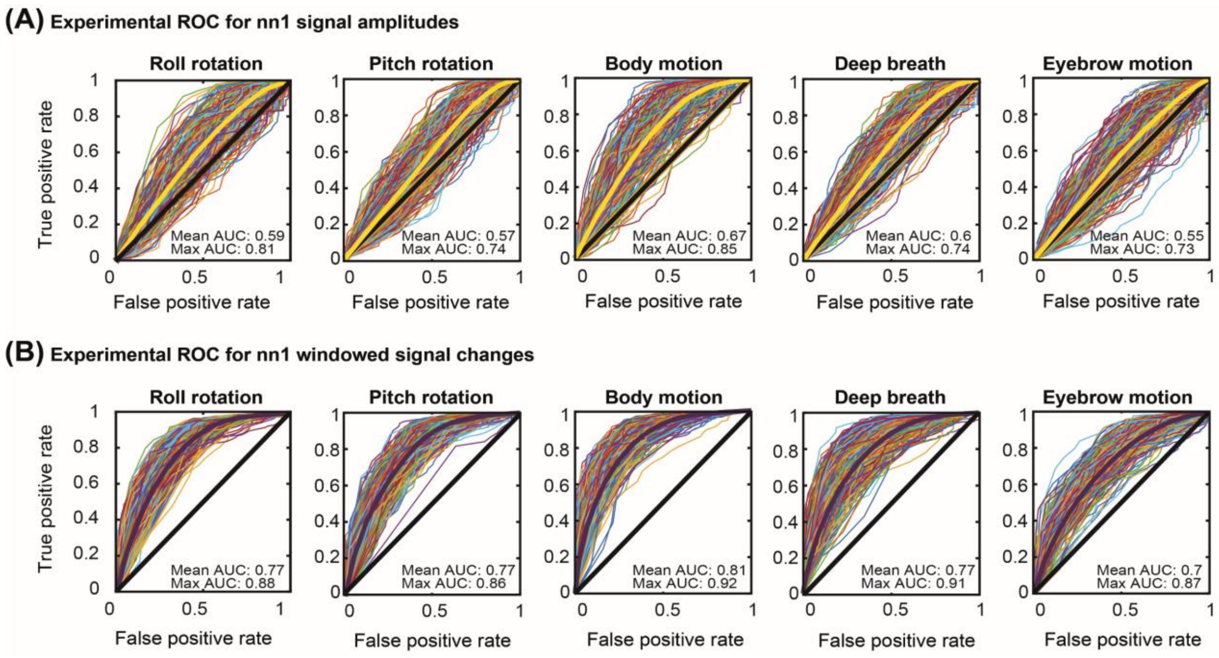


**Figure S3.** The experimental ROC curves are drawn for (A) the absolute signal amplitude of 850 nm nn1 measurements, the yellow curve shows the mean of all ROC curves, and (B) the windowed amplitude changes of the signals in (A), the dark magenta curve shows the mean of all ROC curves. The goal of this figure is to show that regardless of the type of motion, both maximum and mean of the area under the curve (AUC) of the ROC curves in each figure is still lower than or equal to the AUC of GVTD in Fig.4.

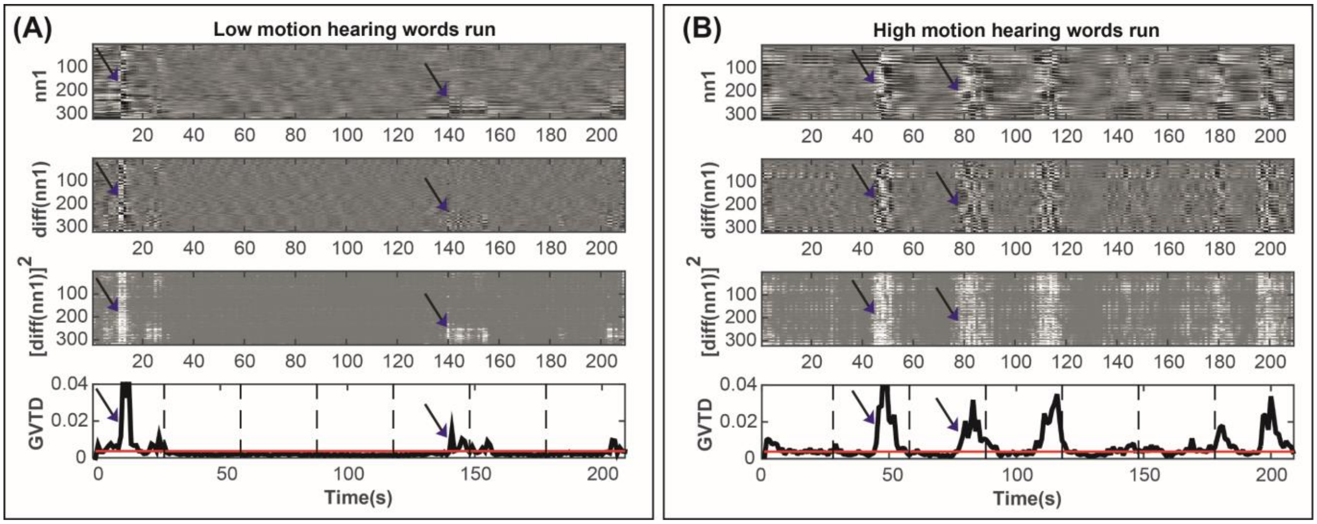


**Figure S4:** GVTD censoring process for the hearing words (HW) task. The four-step process (explained in Fig. 3A-D) for calculating the GVTD time-courses (A) for a low motion subject and (B) a high motion subject in dataset 3. Dashed lines indicate the onsets of the blocks of HW run (6 runs each 30 sec). Red line shows the GVTD threshold calculated based on the mode of the GVTD distribution of each run plus the full length of the left tail (values below mode) of the GVTD histogram. Blocks with time-points exceeding the GVTD threshold are excluded from the analysis (4,5,6 for the low motion subject, and all blocks for the high motion subject). Dark blue arrows indicate some examples of the high contrast in both the gray plots and the spikes of the GVTD time-courses for instances of motion artifacts.

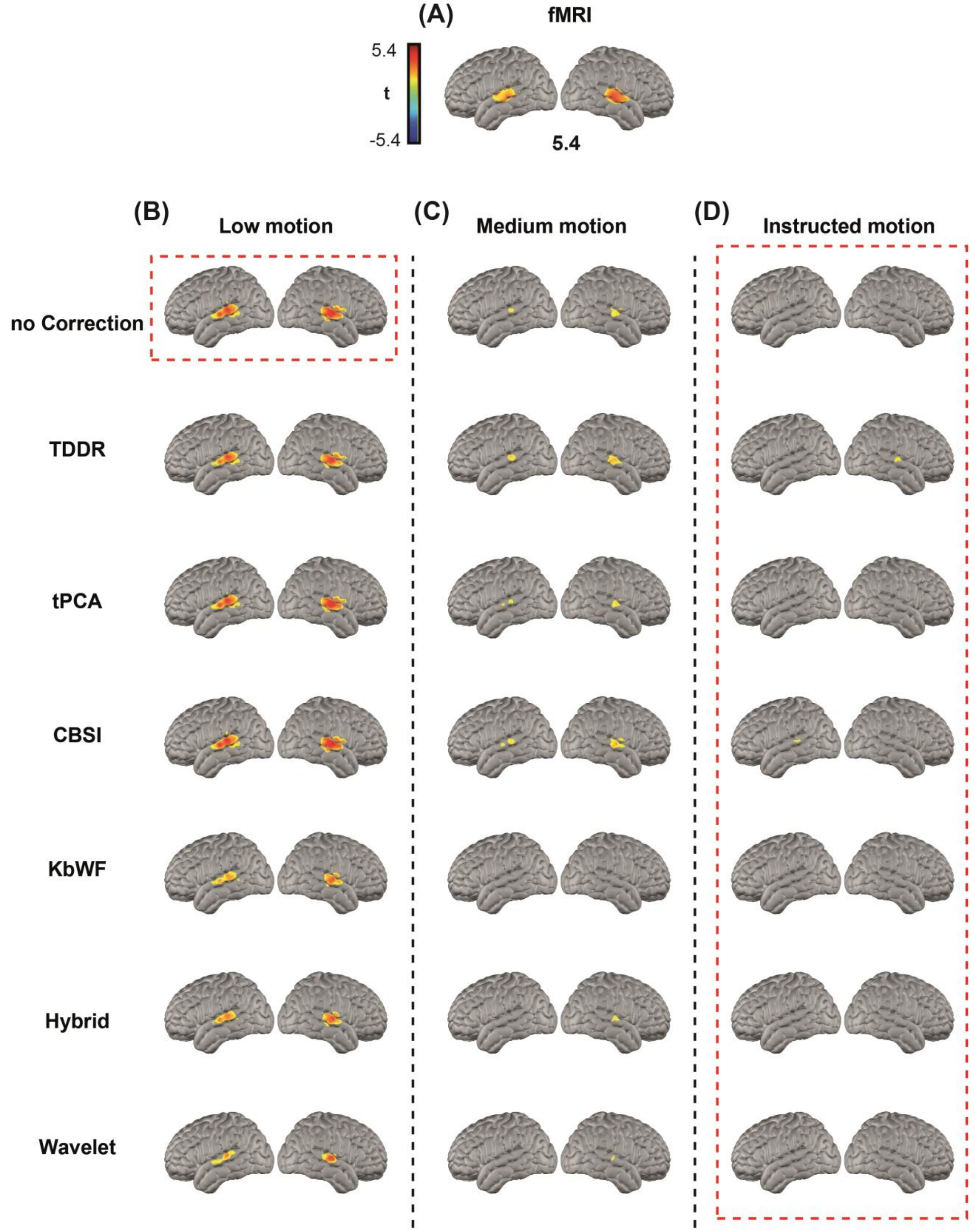


**Figure S5:** HW task t-statistic maps. Voxel-wise maps are shown for (A) Reference fMRI dataset. (B) Low motion data. (C) Medium motion data. (D) Instructed motion data for the three motion level categories determined with mean GVTD scores. Rows represent maps corrected with TDDR, tPCA, CBSI, Kurtosis wavelet, Hybrid, and wavelet filtering methods. All maps are thresholded at $P<0.05$.

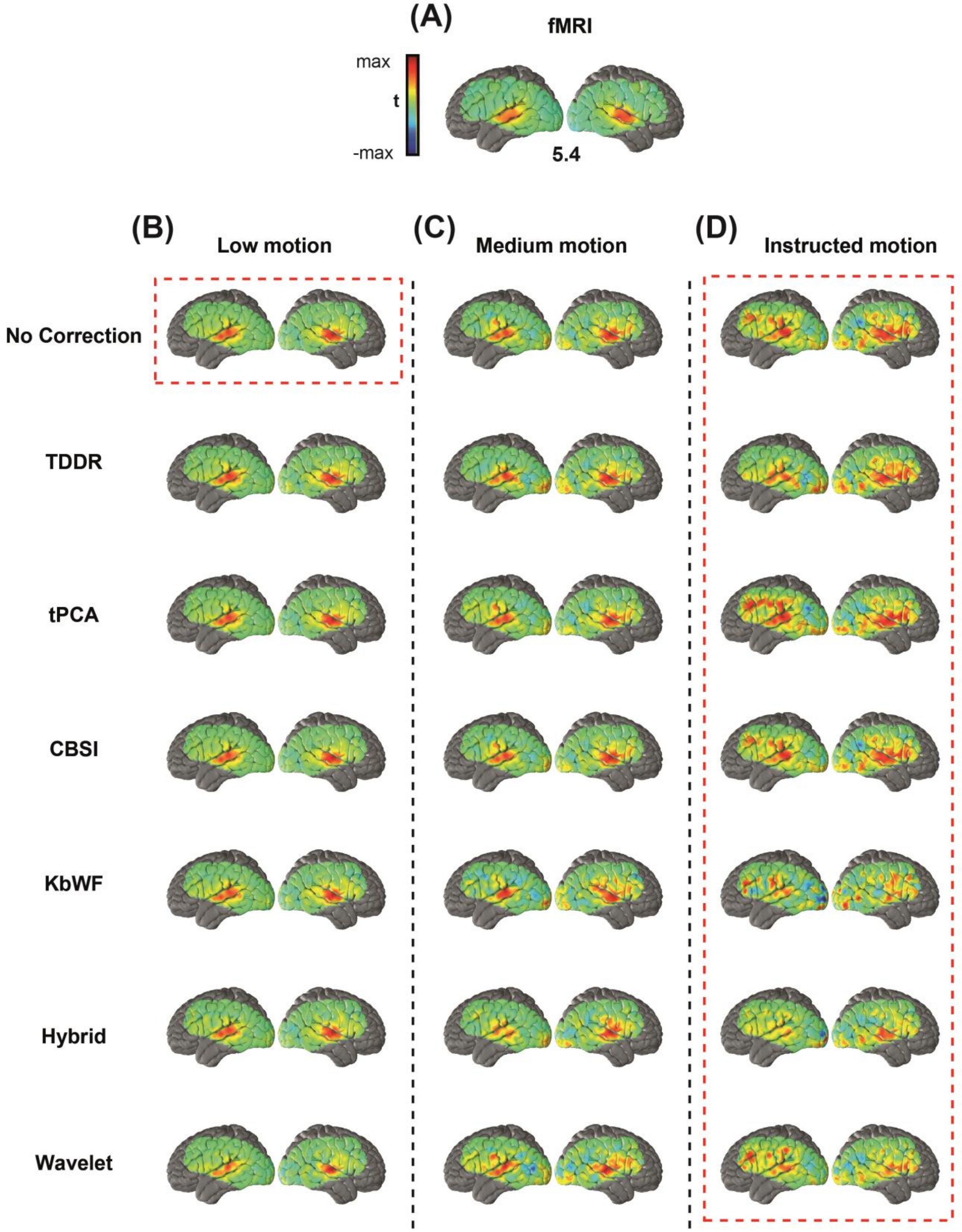


**Figure S6:** Same maps described in Fig. S5 visualized with no thresholding. Voxel-wise maps are shown for (A) Reference fMRI dataset. (B) Low motion data. (C) Medium motion data. (D) Instructed motion data for the three motion level categories determined with mean GVTD scores. Rows represent maps corrected with TDDR, tPCA, CBSI, Kurtosis wavelet, Hybrid, and wavelet filtering methods.
