## Supplementary Tables for "Global Motion Detection and Censoring in High-Density Diffuse Optical Tomography"

| | StatType | numStd | tMotion | tMask | STDEV thresh | AMP thresh | nSV | iqr | kurt | p | FrameSize _sec | tune | filter_cutoff |
|---|---|---|---|---|---|---|---|---|---|---|---|---|---|
| GVTD | Histogram _Mode | 3 | | | | | | | | | | | |
| CBSI | | | | | | | | | | | | | |
| TPCA | | | 0.5 | 2 | 20 | 0.5 | 0.97 | | | | | | |
| Wavelet | | | | | | | | 1.5 | | | | | |
| KbWF | | | | | | | | | 3.3 | | | | |
| Hybrid (S+SG) | | | | | | | | | | 0.99 | 15 | | |
| TDDR | | | | | | | | | | | | 4.695 | 0.5 |

**Table S1:** List of the parameters for motion correction in task data.

| | StatType | numStd | tMotion | tMask | STDEV thresh | AMP thresh | nSV | iqr | kurt | p | FrameSize _sec | tune | filter_cutoff |
|---|---|---|---|---|---|---|---|---|---|---|---|---|---|
| GVTD | Histogram _Mode | 10 | | | | | | | | | | | |
| CBSI | | | | | | | | | | | | | |
| TPCA | | | 0.5 | 2 | 20 | 0.5 | 0.97 | | | | | | |
| Wavelet | | | | | | | | 1.5 | | | | | |
| KbWF | | | | | | | | | 3.3 | | | | |
| Hybrid (S+SG) | | | | | | | | | | 0.99 | 15 | | |
| TDDR | | | | | | | | | | | | 4.695 | 0.5 |

**Table S2:** List of the parameters for motion correction in resting state data.
